## Supplemental figures and tables for "3D RNA-scaffolded wireframe origami"

### Supplementary Information

Molly F. Parsons[1§], Matthew F. Allan[1-3§], Shanshan Li[4,†§], Tyson R. Shepherd[1], Sakul Ratanalert[1,5,‡], Kaiming Zhang[4,†], Krista M. Pullen[1], Wah Chiu[4,6], Silvi Rouskin[2], Mark Bathe[1*]

[1]Department of Biological Engineering, Massachusetts Institute of Technology, Cambridge, MA 02139, United States
[2]Department of Microbiology, Harvard Medical School, Boston, MA, United States 02115
[3]Computational and Systems Biology, Massachusetts Institute of Technology, Cambridge, MA 02139, United States
[4]Department of Bioengineering, Stanford University, Stanford, CA 94305, United States
[5]Department of Chemical Engineering, Massachusetts Institute of Technology, Cambridge, MA 02139, United States
[6]CryoEM and Bioimaging Division, Stanford Synchrotron Radiation Lightsource, SLAC National Accelerator Laboratory, Stanford University, Menlo Park, CA 94025, United States

[†]*Present address*: MOE Key Laboratory for Cellular Dynamics and Division of Life Sciences and Medicine, University of Science and Technology of China, Hefei 230027, China
[‡]*Present address:* Department of Chemical and Biomolecular Engineering, Johns Hopkins University, Baltimore, MD 21218, United States

[§]These authors contributed equally to this work.

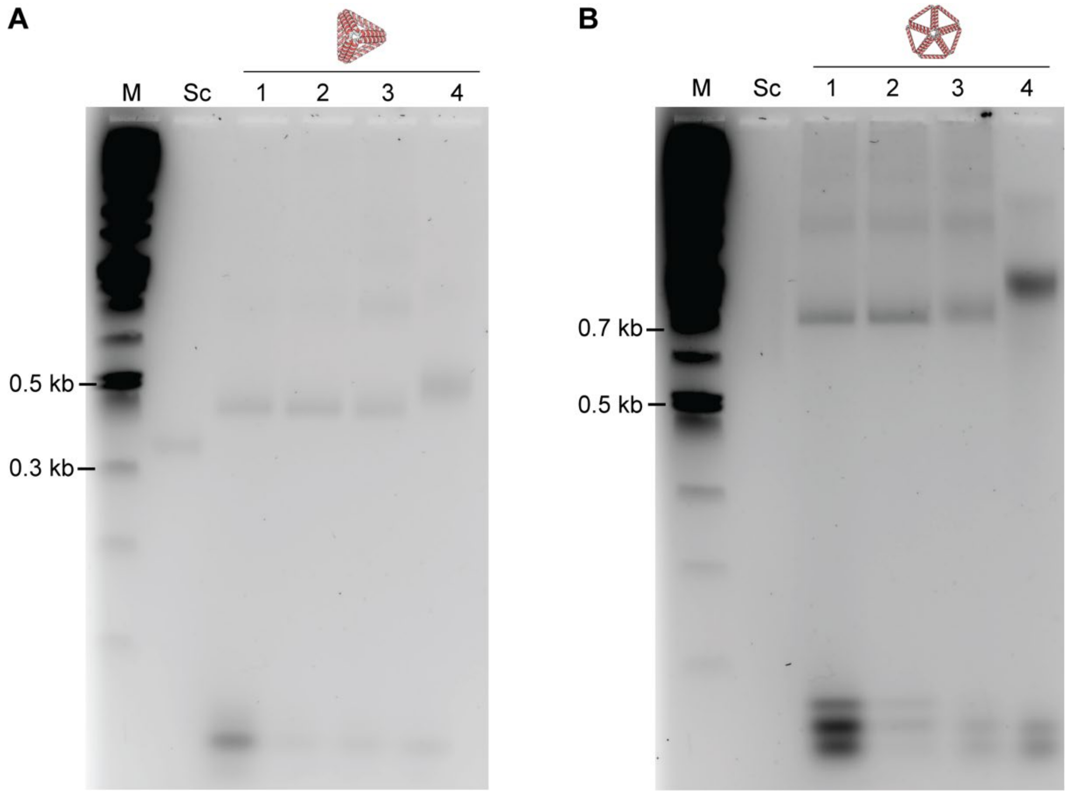

**Figure S1.** Comparison of protocols for folding RNA-scaffolded 3D DX wireframe origami. Protocols compared were 20 nM scaffold in **1.** 10 mM HEPES pH 7.5 with 300 mM KCl, annealed using an overnight folding ramp (adapted from that used for analogous B-form DX wireframe origami[1]) with 20x molar excess of staples; **2.** Same as previous but with 10x staple excess; **3.** 40 mM Tris pH 8.0 with 20 mM acetic acid, 2 mM EDTA, and 12.5 mM magnesium acetate, annealed using a 40 min folding ramp with 10x molar excess of staples (protocol from Wang et al[2]); and **4.** 5 mM Tris pH 7.5 with 1 mM EDTA and 40 mM NaCl, annealed using an overnight folding ramp with 10x molar excess of staples (protocol from Zhou et al[3]). **(A)** Gel mobility shift assay of an EGFP mRNA-scaffolded tetrahedron with six helical turns (66 bp) per edge. **(B)** Gel mobility shift assay of a 23s rRNA-scaffolded pentagonal bipyramid with six helical turns (66 bp) per edge. The “Sc” lanes show the respective scaffold RNA not folded with staples. The high degree of internal structure in the 23s rRNA scaffold often results in a diffuse band in these non-denaturing agarose gels containing 2 mM magnesium acetate. “M” refers to the 1 kb Plus DNA Ladder (NEB) used as a marker for molecular weight.

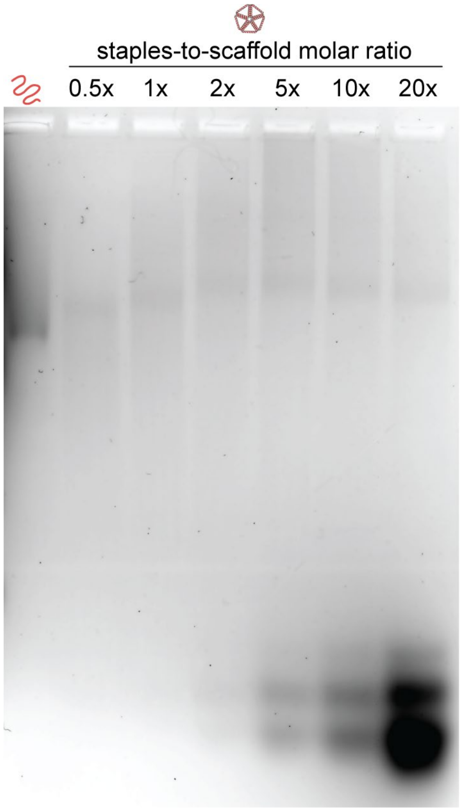

**Figure S2**. Gel mobility shift assay for A-form rPB66 folded with varying molar ratios of staples-to-scaffold. The upward shift of the folded rPB66 band appears to stabilize at approximately 2x staples-to-scaffold ratio, suggesting this is the minimum amount of staples to use for proper folding. A 1x staples-to-scaffold molar ratio led to a reduced upward shift of the folded band relative to the scaffold band, suggesting only partial folding, or perhaps reduced yield of correct structures similar to observations by Wang et al. when using a less than 1x staples-to-scaffold ratio for 2D RNA:DNA origami. The reduced staples-to-scaffold ratio requirement for RNA:DNA origami relative to DNA:DNA origami is likely due to the higher affinity in the hybrid duplexes of the former[2].

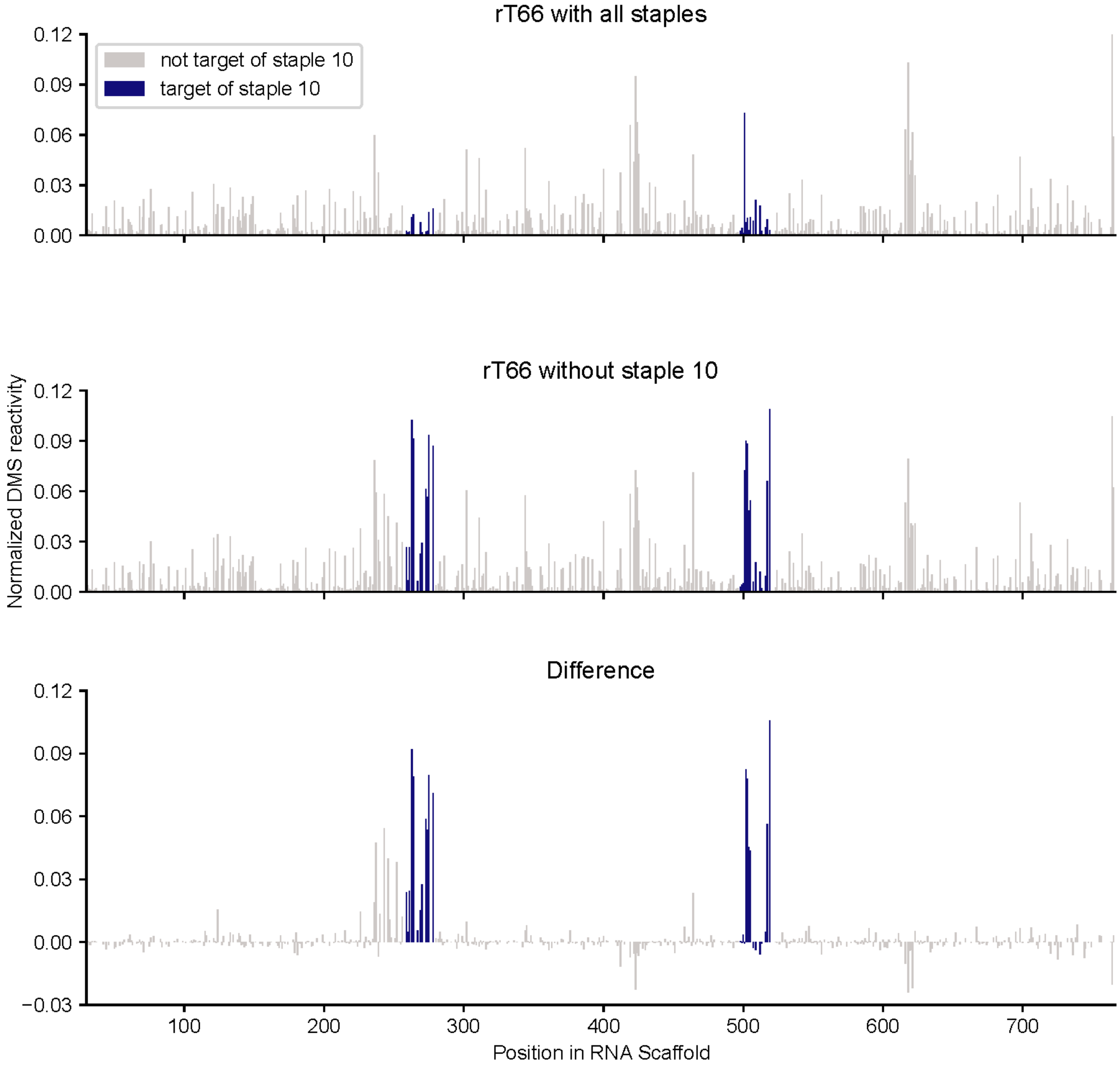

**Figure S3**. DMS-MaPseq reactivity profiles for the A-form rT66 scaffolded with EGFP mRNA, folded with and without staple 10, whose targeted nucleotides are shown in blue.

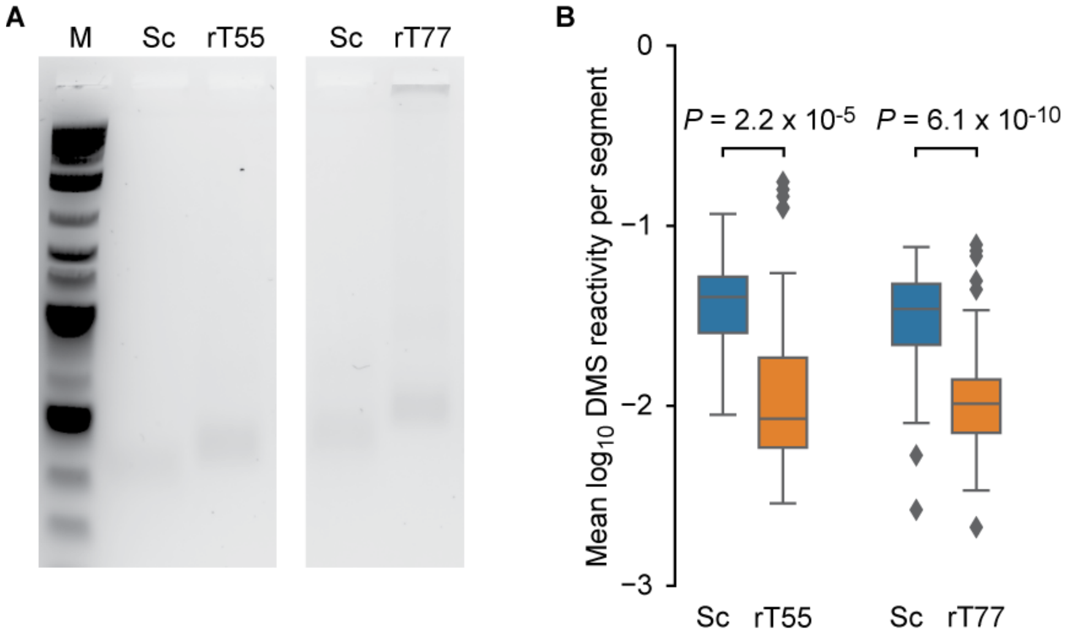

**Figure S4**. Gel mobility shift assays for A-form RNA-scaffolded tetrahedron with odd edge lengths. **(A)** (left) Tetrahedron with five helical turns per edge, with 'rsc1218v1_T55' synthetic RNA fragment scaffold and (right) tetrahedron with seven helical turns per edge, with 'rsc1218v1_T77' synthetic RNA fragment scaffold. M indicates the 1kb plus DNA ladder (NEB) used as a marker. Sc indicates unfolded scaffold. **(B)** Box plots of normalized DMS reactivity per double-helical segment for the two odd-edge-length tetrahedral and their scaffolds.

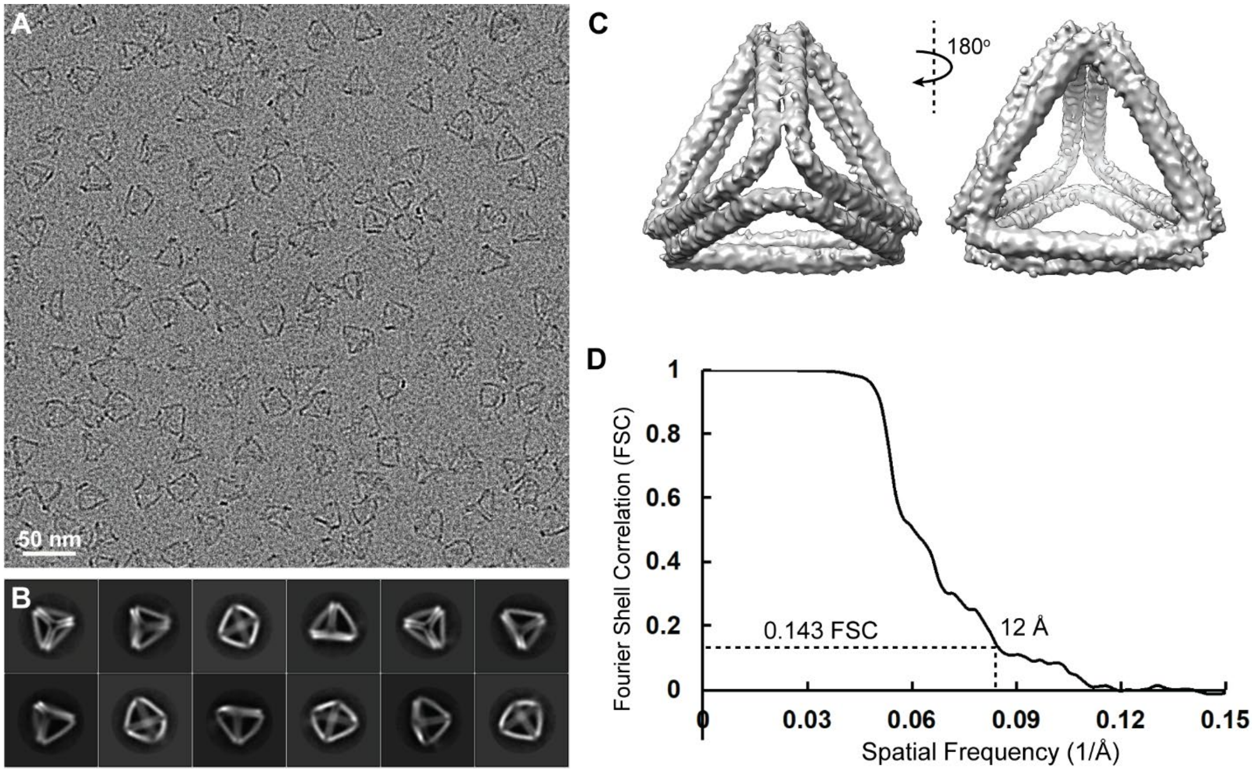

**Figure S5**. Cryo-EM reconstruction for the A-form tetrahedron with EGFP mRNA scaffold (rT66). **(A)** Representative micrograph. **(B)** 2D class averages. **(C)** Two views of the reconstruction. **(D)** Fourier shell correlation plot; the resolution of the reconstruction is 12Å.

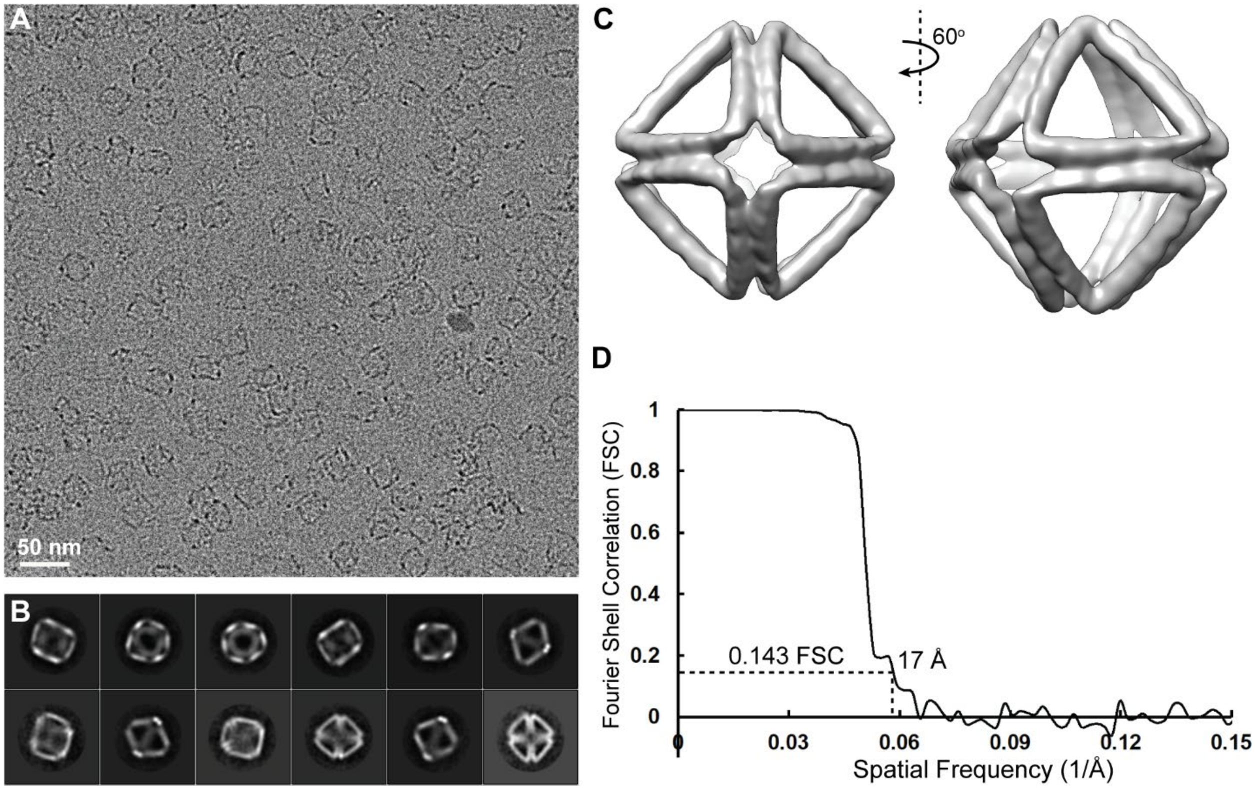

**Figure S6**. Cryo-EM reconstruction for the A-form octahedron with four helical turns per edge with M13 transcript scaffold (rO44). **(A)** Representative micrograph. **(B)** 2D class averages. **(C)** Two views of the reconstruction. **(D)** Fourier shell correlation plot; the resolution of the reconstruction is 17Å.

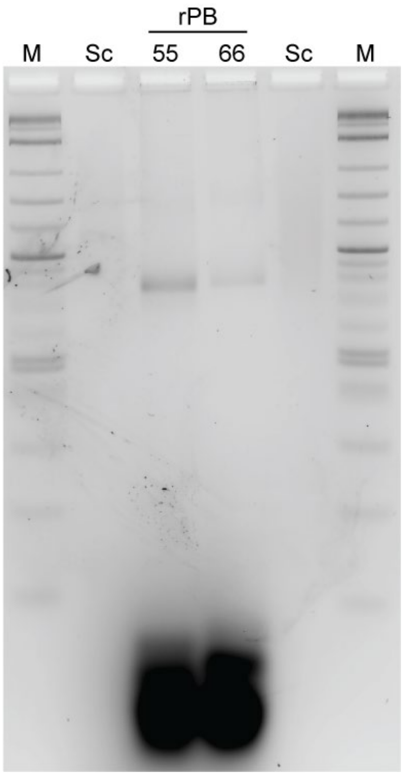

**Figure S7**. Gel mobility shift assay for A-form pentagonal bipyramid with five helical turns per edge (rPB55), with 23s rRNA fragment scaffold. This scaffold often forms a smear in agarose gels with salt, likely due to the formation of secondary structure, but it shows a single band on denaturing PAGE gels. The pentagonal bipyramid with six helical turns per edge (rPB66) folded with the same scaffold is included in the gel for comparison.

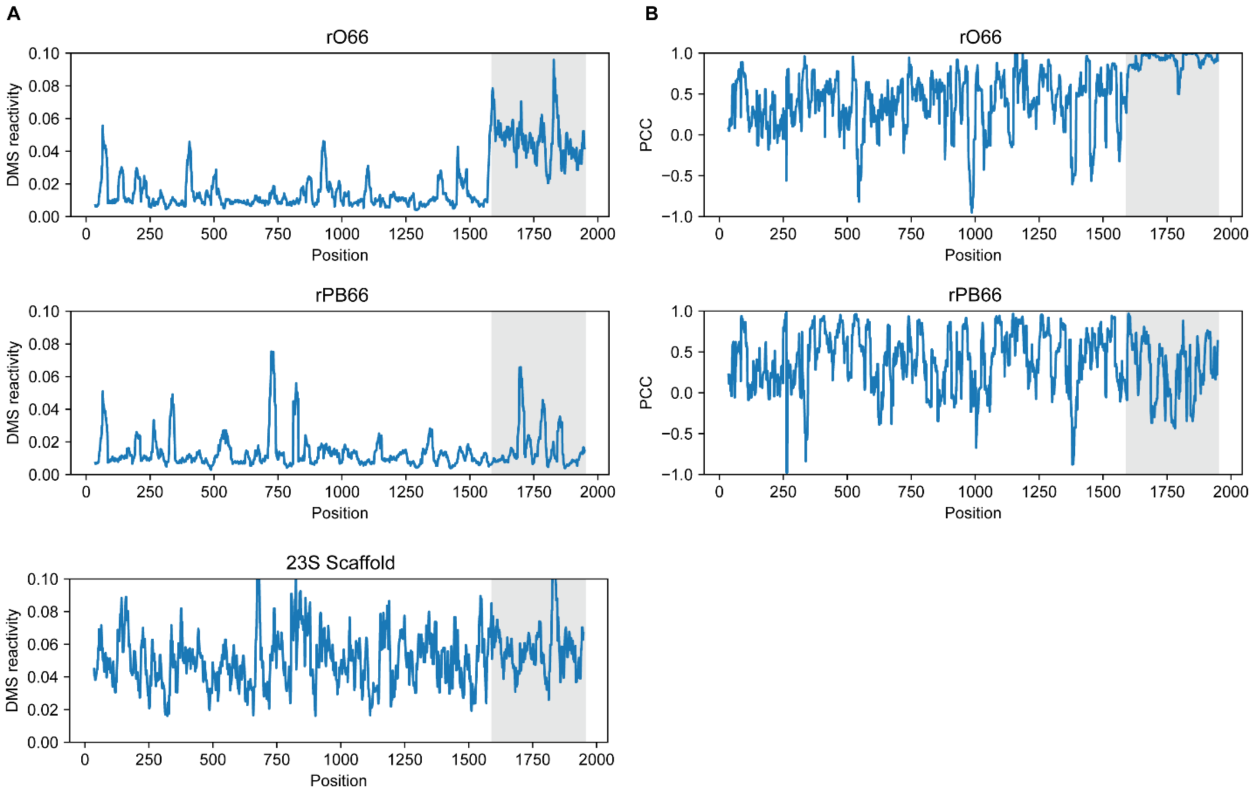

**Figure S8**. DMS-MaPseq reactivity profiles for A-form 23s-rRNA-scaffolded origami. **(A)** Normalized DMS reactivity for each nucleotide of the scaffold, when incubated with staples to fold the rO66 or the rPB66, or when incubated without staples (bottom). The final 396 nt of the scaffold (highlighted in grey) were not targeted by staples in the rO66. **(B)** Pearson Correlation Coefficient (PCC) values for the DMS reactivities of the scaffold when folded with staples vs. the scaffold folded alone. The final 396 nt of the scaffold (highlighted in grey) were not targeted by staples in the rO66.

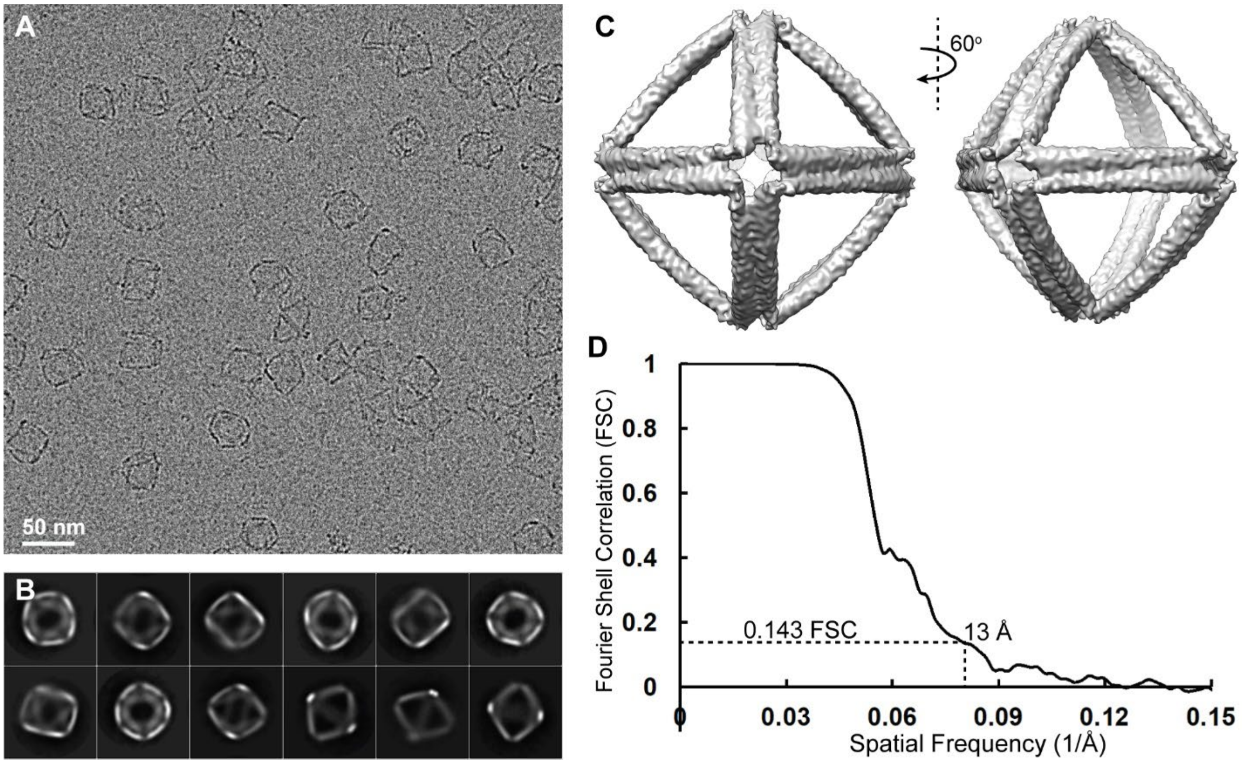

**Figure S9**. Cryo-EM micrographs for the A-form octahedron with six helical turns per edge and 23s rRNA fragment scaffold (rO66). **(A)** Representative micrograph. **(B)** 2D class averages. **(C)** Two views of the reconstruction. **(D)** Fourier shell correlation plot; the resolution of the reconstruction is 13Å.

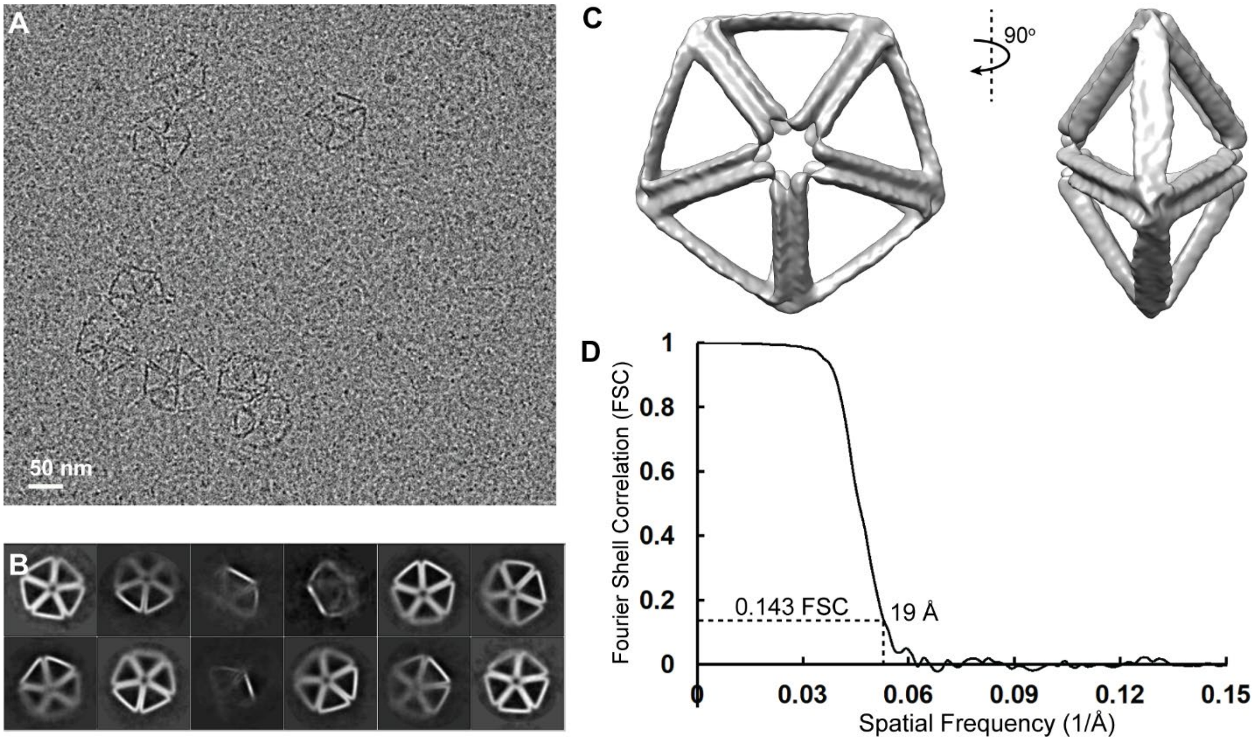

**Figure S10**. Cryo-EM reconstruction for the A-form pentagonal bipyramid with 23s rRNA fragment scaffold (rPB66). **(A)** Representative micrograph. **(B)** 2D class averages. **(C)** Two views of the reconstruction. **(D)** Fourier shell correlation plot; the resolution of the reconstruction is 19Å.

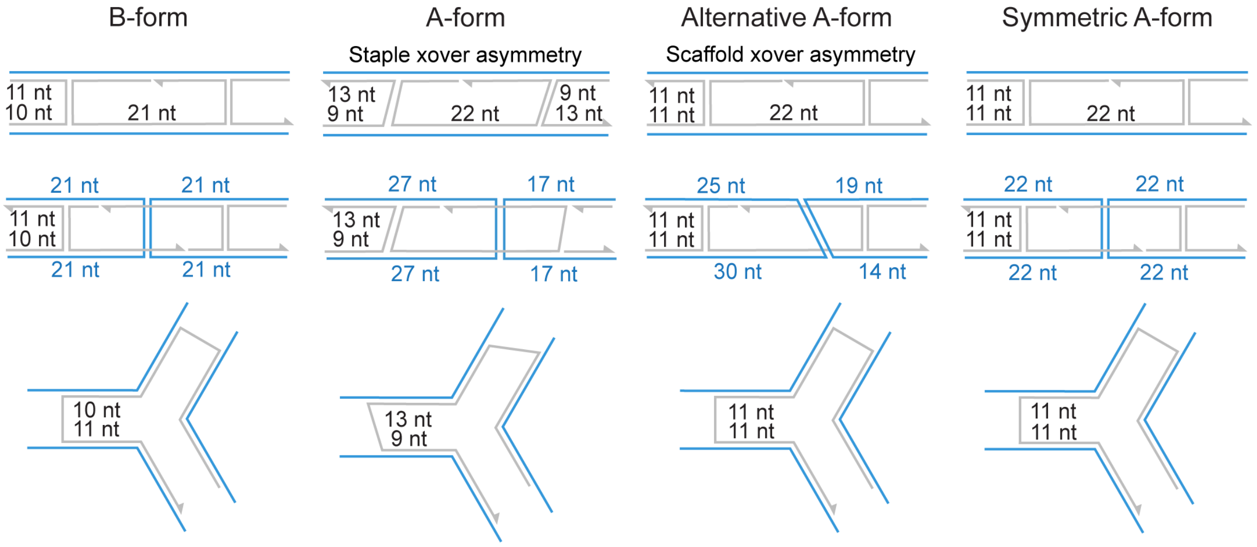

**Figure S11**. Comparison of B-form (DAEDALUS[1]), A-form, Alternative A-form (Alt A-form), and Symmetric A-form (Sym A-form) scaffold and staple edge and vertex routings. "xover" denotes "crossover."

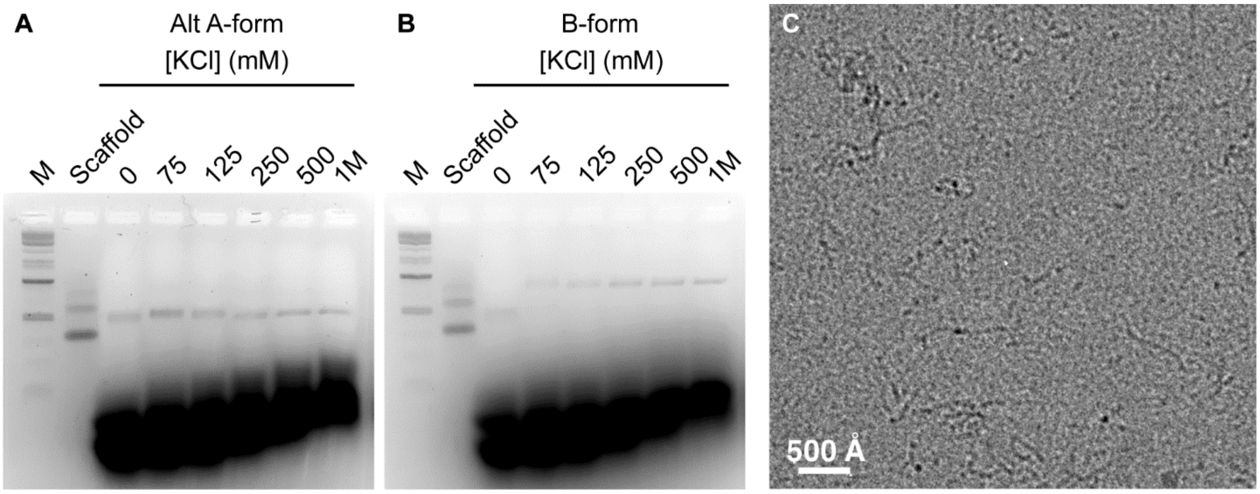

**Figure S12**. B-form staple designs from DAEDALUS[1] used to fold EGFP mRNA-scaffolded origami. **(A)** KCl titration of Alt A-form staple designs with RNA scaffolding, designed to fold a tetrahedron with six helical turns (66 bp) per edge. **(B)** KCl titration of B-form staple designs with RNA scaffolding, designed to fold a tetrahedron with six helical turns (63 bp) per edge, with the band being noticeably higher, suggesting a less compact (less complete) folded product. **(C)** Cryo-electron microscopy micrograph showing the unfolded structure of the B-form-stapled origami.

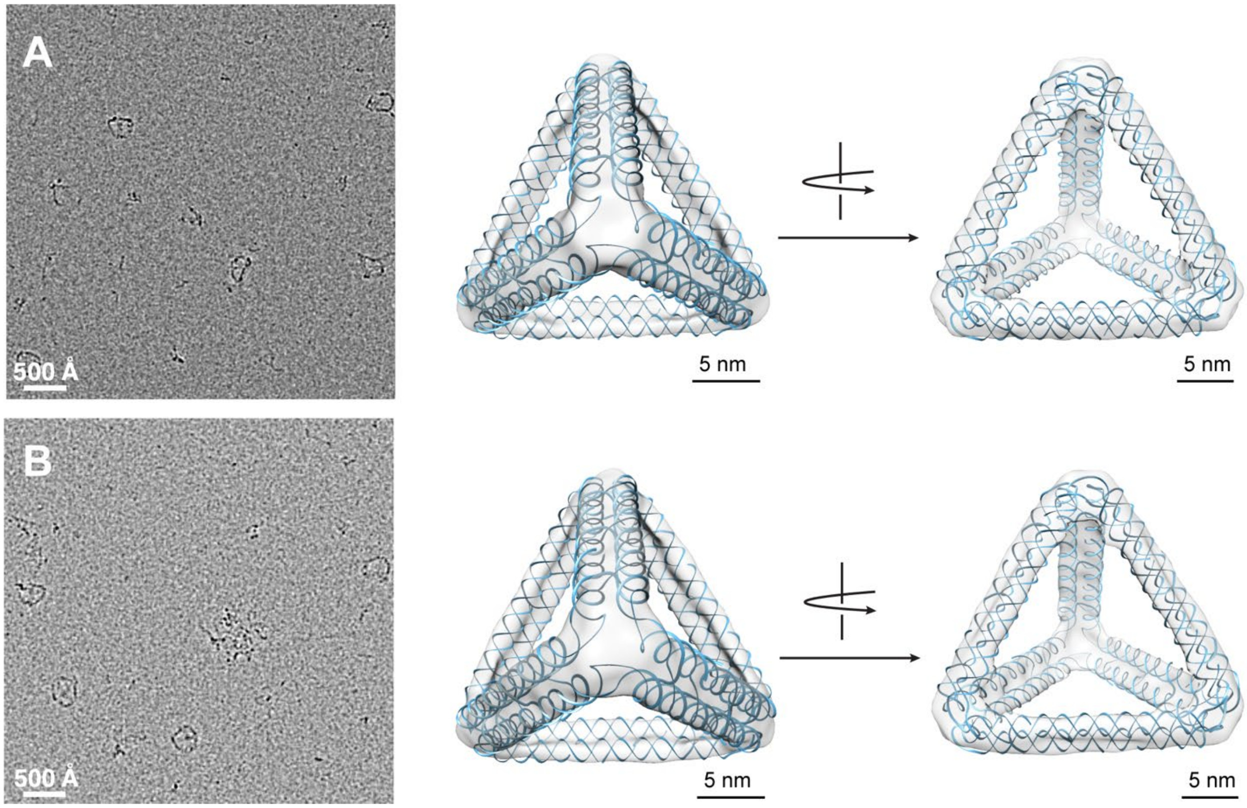

**Figure S13**. Cryo-EM comparison of RNA-scaffolded tetrahedron with 66-bp edge lengths folded using (**A**) Alt A-form geometry staples or (**B**) Sym A-form geometry staples.

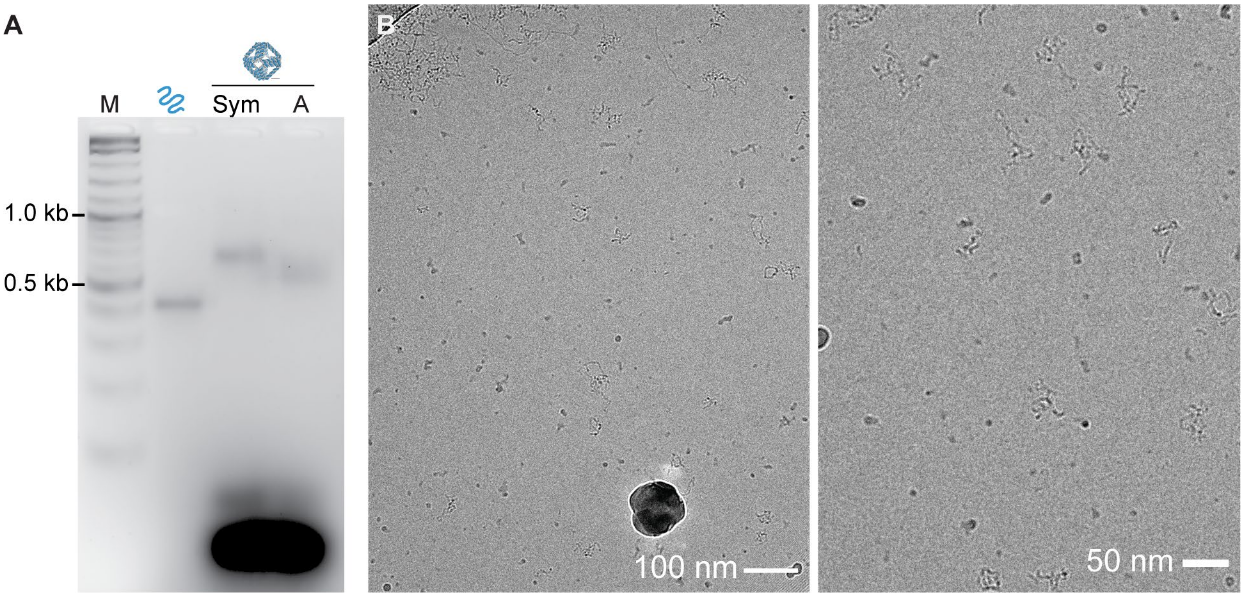

**Figure S14**. Cryo-EM micrographs for a Sym A-form regular octahedron with four helical turns (44 bp) per edge, scaffolded with M13 transcript RNA. **(A)** Gel mobility shift assay comparing the Sym A-form and A-form octahedra folds to the unfolded scaffold. The Sym A-form fold leads to a notably higher band shift than the A-form fold. The marker (M) is 1kb plus DNA ladder from NEB. **(C)** Cryo-EM micrographs of the Sym A-form octahedron, showing no well-folded octahedral particles.

For an alternative A-form (Alt A-form) EGFP mRNA-scaffolded tetrahedron with six helical turns (66 bp) per edge, gel mobility shift assays after folding with KCl or NaCl showed an upward shift of the major band relative to unpaired scaffold, with the band position and breadth stabilized at 300 mM monovalent salt (**Supplementary Fig. S15**). These results suggested RNA:DNA origami wireframe particles were properly folded in 300 mM monovalent salt and 10 mM HEPES-KOH pH 7.5. No major band was observed when attempting to fold the tetrahedron in magnesium, as expected, likely due to RNA degradation from this divalent salt at elevated temperatures during annealing (**Supplementary Fig. S16**). Consistent with this hypothesis, this effect was mitigated by using a fast-folding protocol[2] (**Supplementary Fig. S17**). Higher yields of the folded particles were achieved in HEPES-KOH pH 7.5 buffer than in Tris-HCl pH 8.1 buffer (**Supplementary Fig. S17**), possibly due to the combined effects of higher pH and temperature.

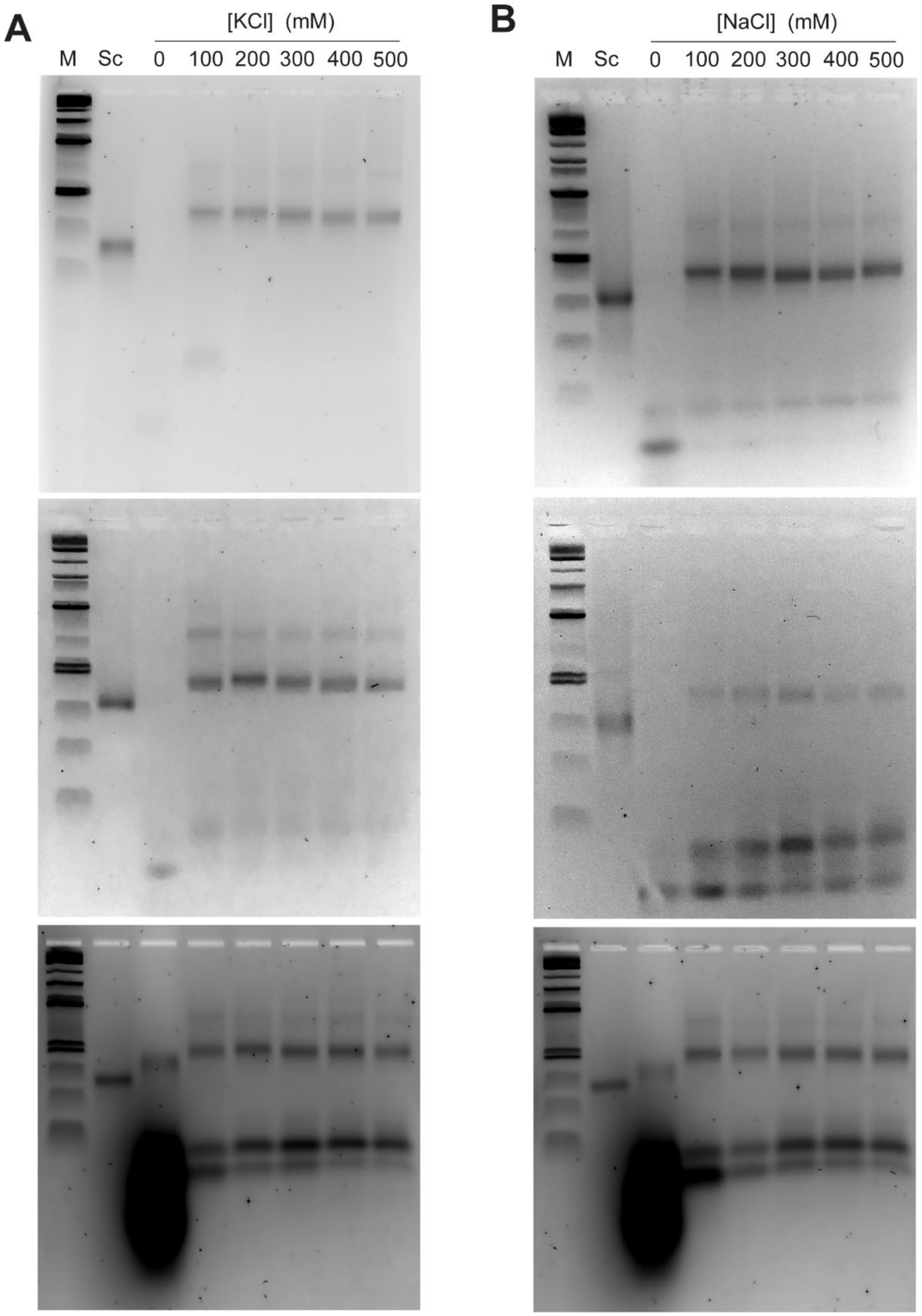

**Figure S15.** Triplicate titration of folding in monovalent salts and 10 mM HEPES-KOH pH 7.5 for the Alt A-form rT66. **(A)** Titration series of KCl. **(B)** Titration series of NaCl. M is 1kb plus DNA ladder from New England Biosciences. Sc indicates the scaffold alone. The folded origami bands shift slightly up relative to the scaffold band, with mobility and breadth stabilizing at 300 mM salt.

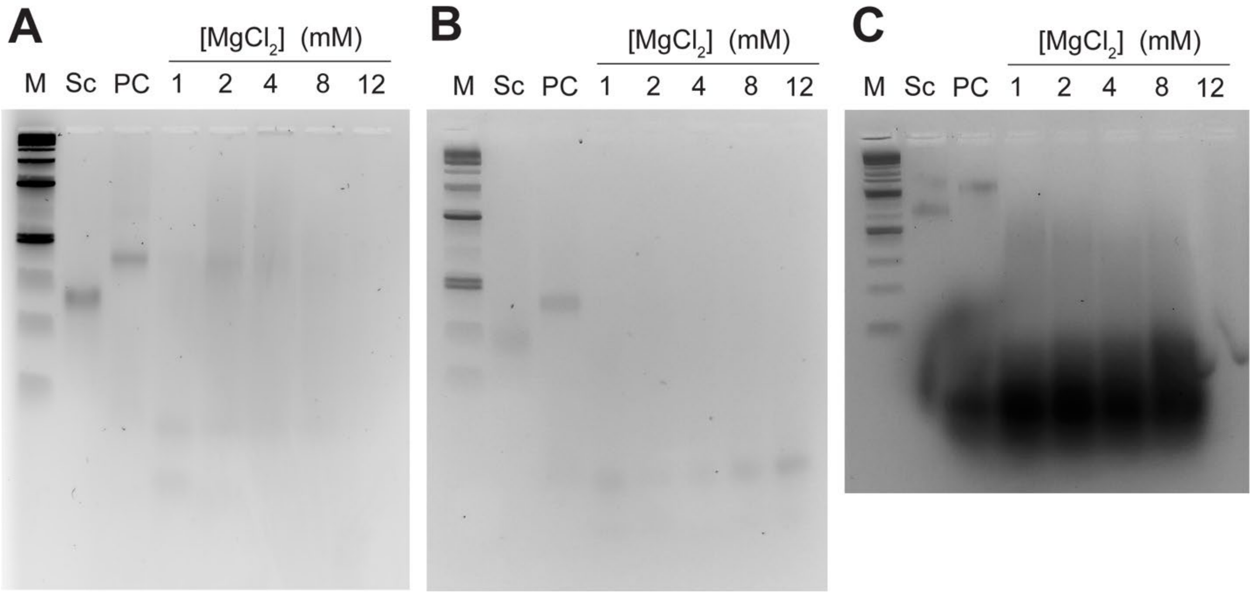

**Figure S16. (A)** and **(B)** Duplicate titration of folding in $MgCl_2$ and 10 mM HEPES-KOH pH 7.5 for the Alt A-form rT66. **(C)** A titration of folding in $MgCl_2$ for the Alt A-form rPB66. M is 1kb plus DNA ladder from New England Biosciences. Sc indicates the scaffold alone. PC indicates a positive control of the origami folded in 300 mM KCl and 10 mM HEPES-KOH pH 7.5. In all cases, no discrete band corresponding to the folded origami or scaffold appears at any concentration of magnesium, suggesting at least partial degradation of the RNA scaffold.

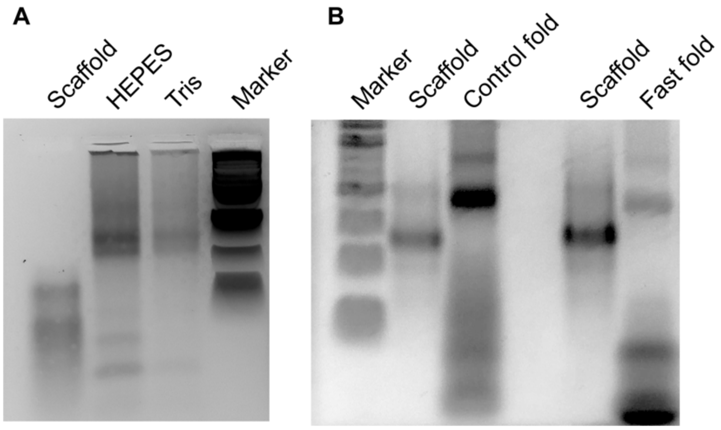

**Figure S17**. Comparison of alternative folding conditions for the Alt A-form rT66. (**A**) HEPES pH 7.5 was compared to Tris-HCl pH 8.1 for the 13-h folding protocol modified from DAEDALUS[1], both with KCl. (**B**) HEPES pH 7.5 and KCl with the 13-h folding protocol ("Control fold") was tested against a previously published 40-min protocol in TAE buffer with magnesium ("Fast fold"[2]), showing nearly equivalent yields when adjusted for loading amounts.

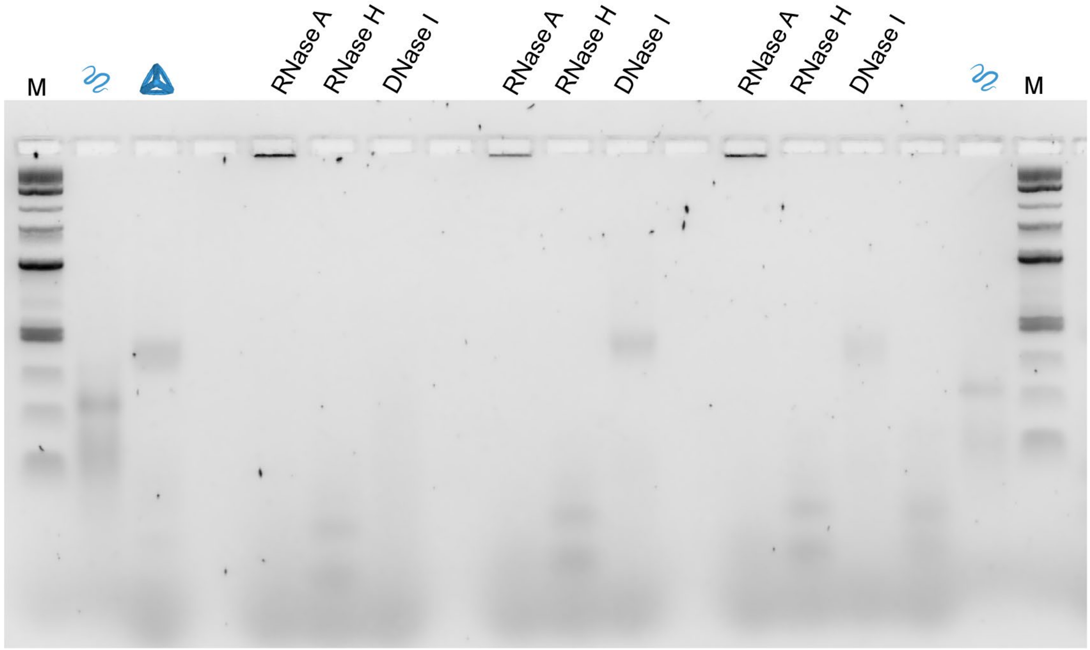

**Figure S18.** Triplicate characterization of biochemical stability of the Alt A-form EGFP mRNA-scaffolded tetrahedron with 66-bp edge length, using canonical nucleotides. The RNase A-, Rnase H-, and Dnase I-labeled lanes all represent the folded rT66 treated with the respective nuclease for 5 min at 37ºC. Two of the three Dnase I replicates show the intact folded origami band. Rnase H, which specifically targets RNA hybridized to DNA, degraded the folded origami and released staples within five minutes, with the folded origami band completely disappearing, the scaffold strand appearing to be fully digested, and bands corresponding to DNA staples concomitantly appearing. This degradation additionally confirmed trace template DNA was not responsible for forming the principal origami product. With Rnase A treatment, no folded origami gel band showed after five minutes incubation, and some aggregation appeared, possibly due to intact origami-bound Rnase A without cleavage because of the steric hindrance in the double-duplex edges. This interpretation is supported by the absence of discrete bands corresponding to DNA staple strands after Rnase A treatment, since they are not digested by Rnase A. Interestingly, Dnase I did not appear to degrade incorporated DNA staples within a 5-minute incubation, suggesting that nanostructuring with RNA might offer some protection to DNA staples from enzymatic degradation.

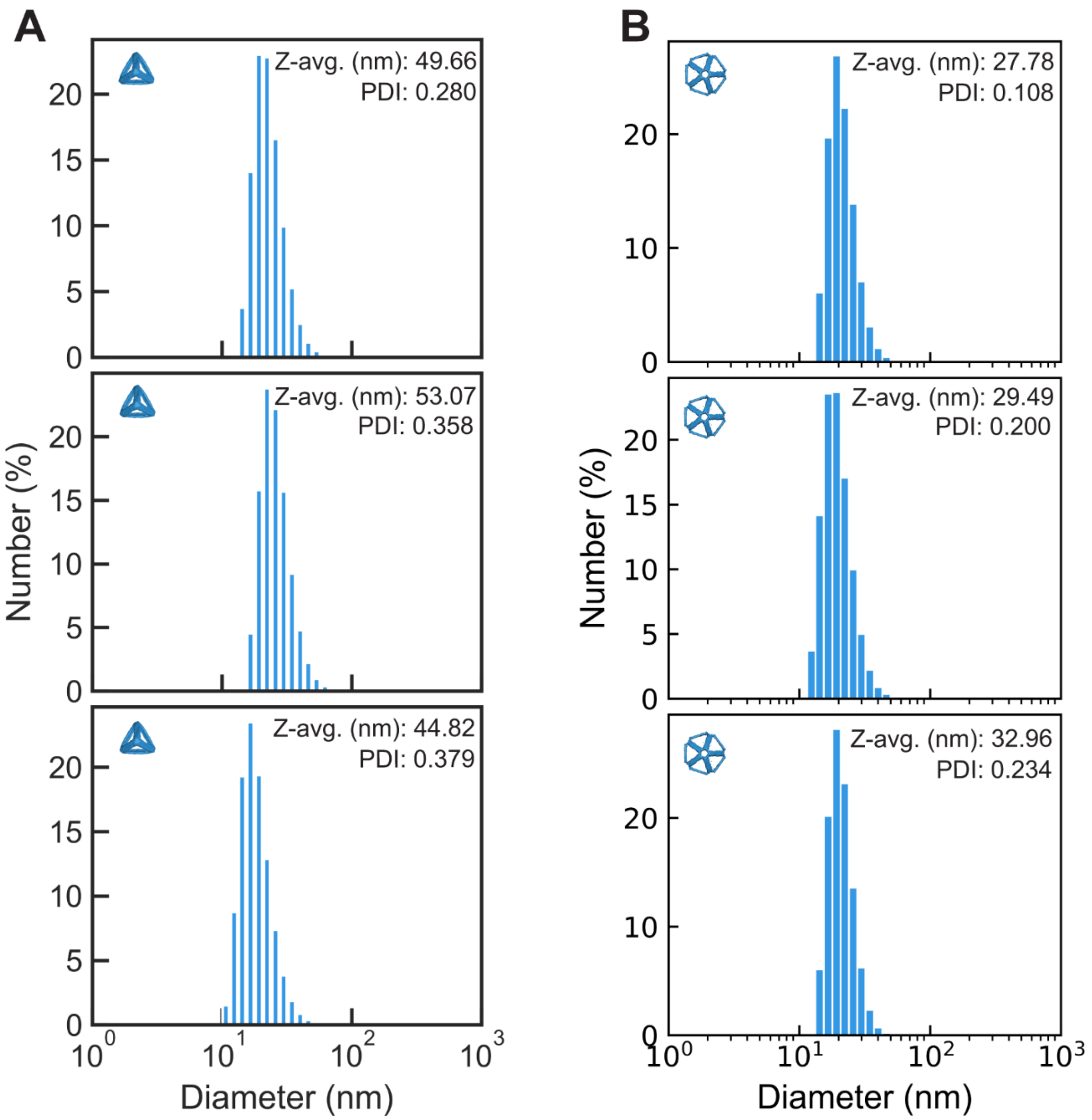

**Figure S19**. Dynamic light scattering (DLS) showing monodisperse objects for **(A)** Alt A-form rT66. The size (hydrodynamic diameter) at the peak of the distribution by number is 17.86 nm ± 5.67 nm. The designed edge length is 17.16 nm and expected tetrahedral height is approximately 14 nm. **(B)** Alt A-form rPB66. The size (hydrodynamic diameter) at the peak of the distribution by number is 20.38 nm ± 5.10 nm. The designed edge length is 17.16 nm and expected height and width of the pentagonal bipyramid are approximately 21 nm x 36 nm.

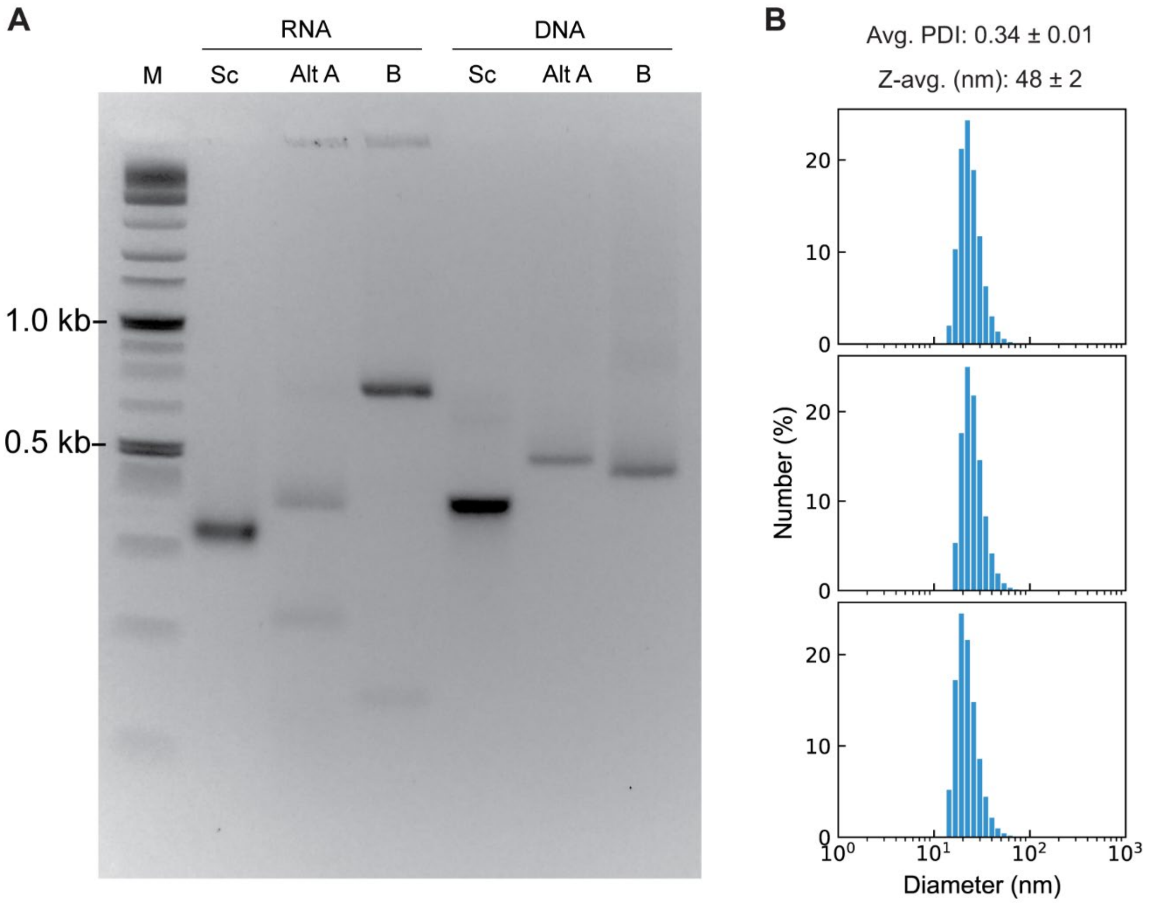

**Figure S20**. Comparison of RNA- and DNA-scaffolded tetrahedra with six helical turns (66 bp for Alt A-form, 63 bp for B-form) per edge. **(A)** Prokaryotic EGFP mRNA sequence was used for either RNA or DNA scaffold (Sc), folded with DNA staples designed for Alt A-form in 10 mM HEPES and 300 mM KCl; or folded with DNA staples designed for B-form routing following the DAEDALUS protocol[1]. All folded objects show discrete bands that are shifted slightly upwards relative to the scaffold band, suggesting proper folding, with the exception of RNA scaffold folded with B-form staples that leads to an extreme upward band shift and appears to not have folded compactly. **(B)** Triplicate dynamic light scattering measurement of the B-form DNA-scaffolded tetrahedron with six helical turns (63 bp) per edge. The size (hydrodynamic diameter) at the peak of the distribution by number is 21.46 nm $\pm$ 6.46 nm. The designed edge length is 21.42 nm and expected height is approximately 17 nm. As expected, the DNA-scaffolded B-form tetrahedron had a slightly larger hydrodynamic diameter than the A-form tetrahedron with the same number of helical turns per edge, corresponding to the greater axial rise per duplex turn in B-form compared with A-form duplex geometries.

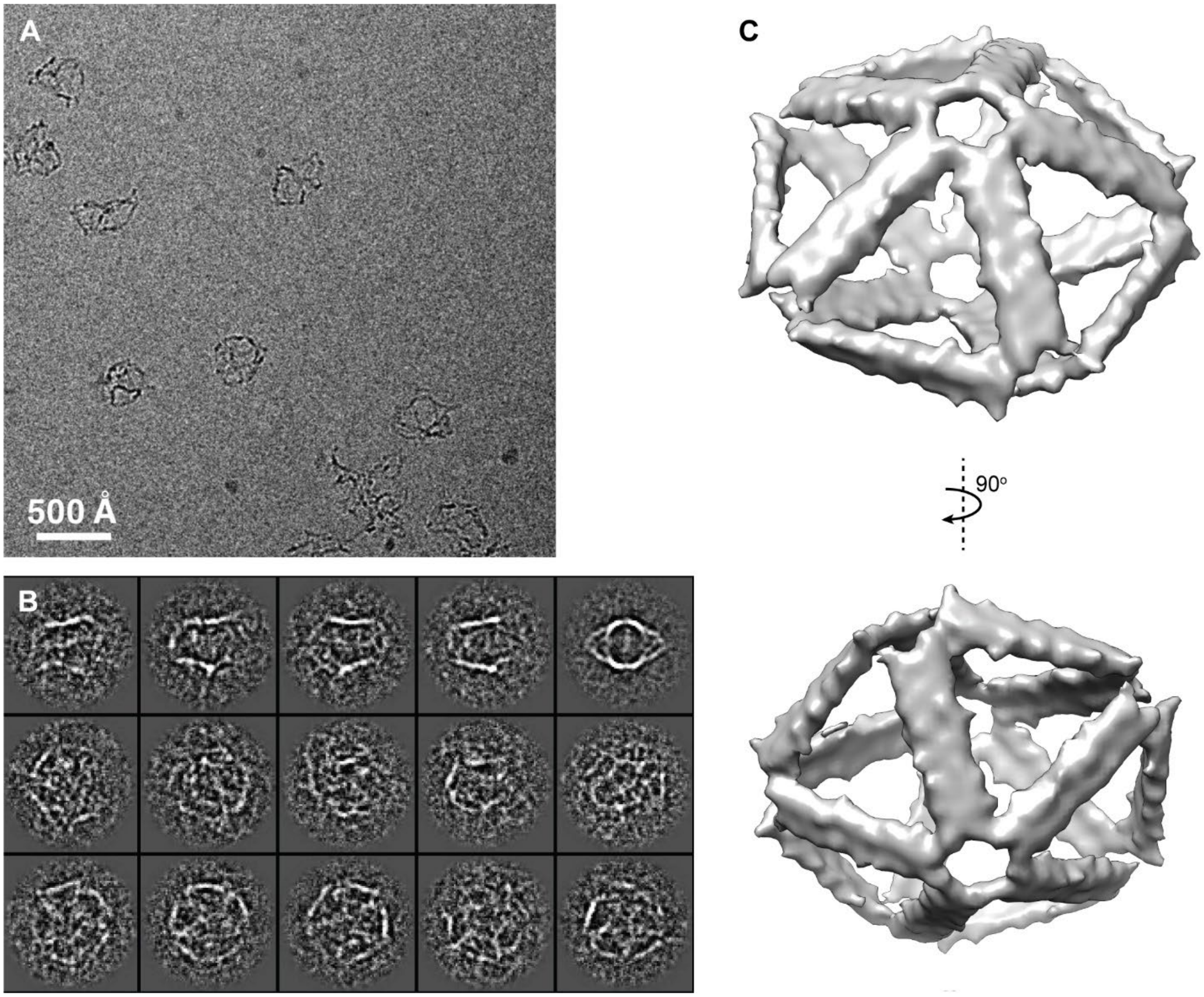

**Figure S21**. Cryo-EM micrograph of Alt A-form RNA-scaffolded pentagonal bipyramid with 66-bp edge lengths. **(A)** Representative micrograph. **(B)** 2D class averages. **(C)** Two orthogonal views of the reconstruction.

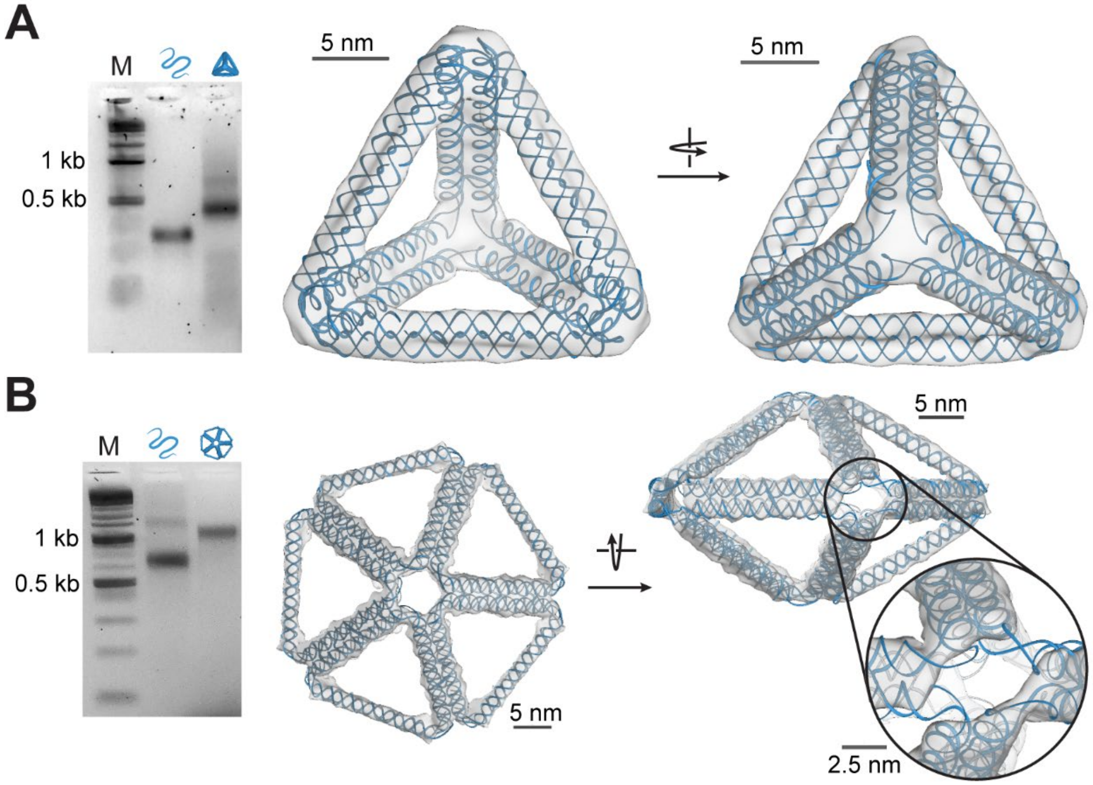

**Figure S22.** 3D structural characterization of Alt A-form RNA-scaffolded wireframe nanoparticles. (**A**) A regular Alt A-form RNA-scaffolded tetrahedron showing the distinct wireframe structure. (**B**) A regular Alt A-form RNA-scaffolded pentagonal bipyramid with 66-bp edge lengths showing the wireframe structure. A notable twist is seen along the edge, which disrupts the electron density at the vertices, as shown.

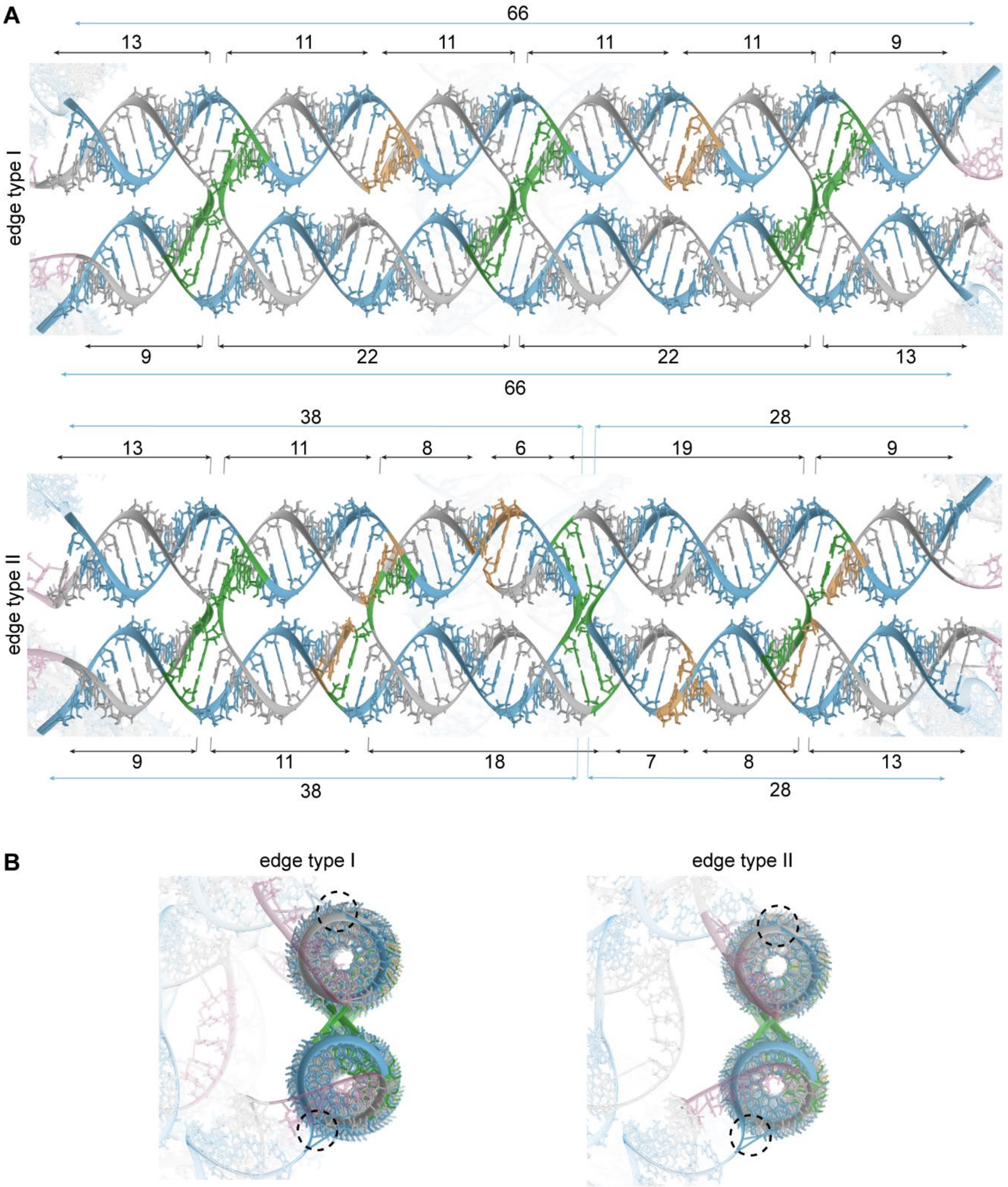

**Figure S23.** Close-up of edges on the output 3D structure predictions for an A-form tetrahedron with six helical turns per edge, showing both an edge with and an edge without a scaffold crossover. **(A)** Top-down views of the full edges. The connectivity and nucleotide counts in the structure predictions matches the intended design (schematized in Figure S11 for four helical turns per edge). Blue: scaffold strand. Grey: staple strands. Green: base pairs at crossovers. Yellow: base pairs at staple nicks. Pink: poly-T loops, bridging to neighboring edges. **(B)** End views of the edges. The atomic model is generated with an assumption that the scaffold (blue) will exit the duplex at a helical position opposite the other duplex of the edge, so that the scaffold is near to its entry point in the subsequent edge and no unpaired scaffold is required.

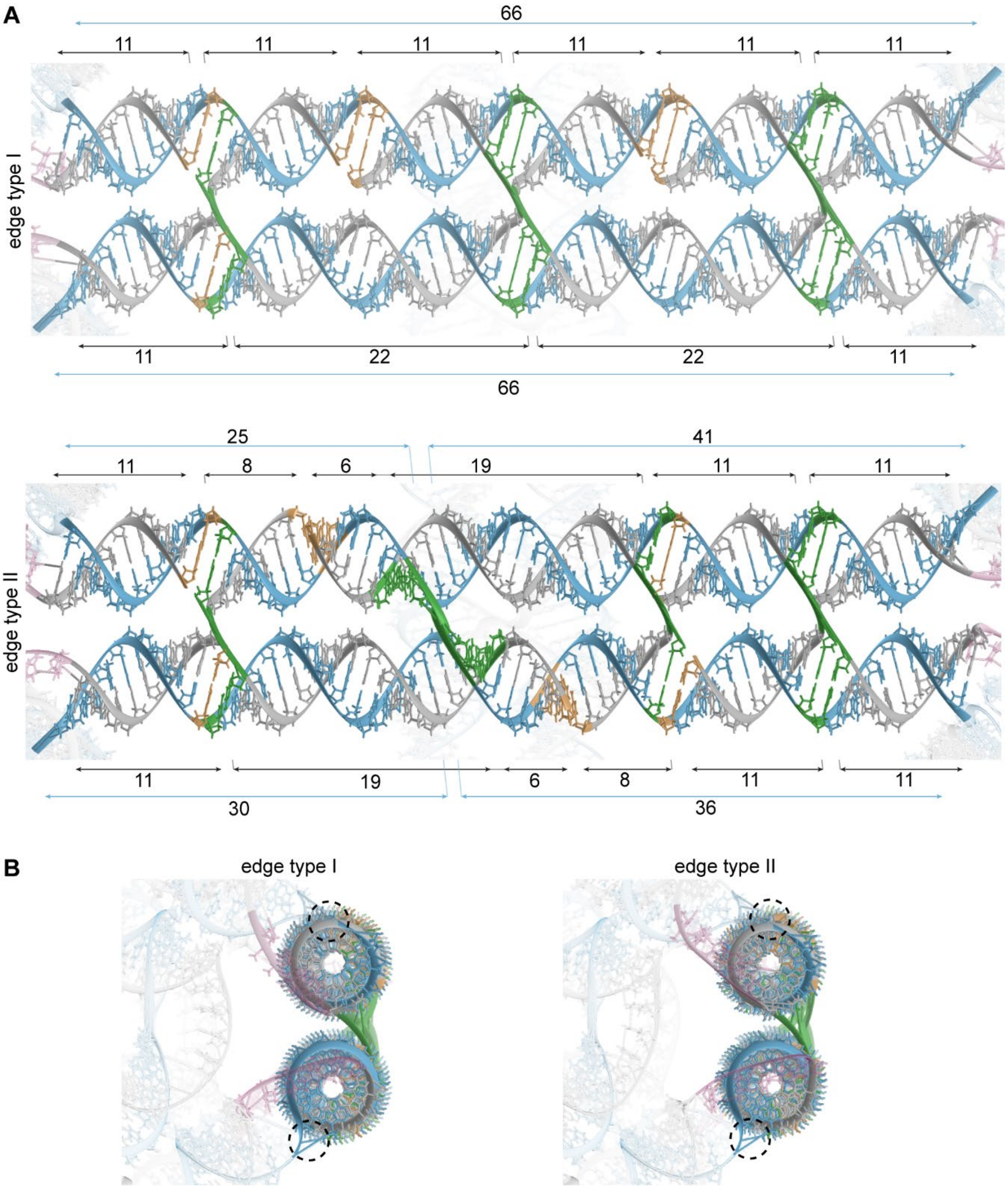

**Figure S24.** Close-up of edges on the output 3D structure predictions for an Alt A-form tetrahedron with six helical turns per edge, showing both an edge without (type I) and an edge with (type II) a scaffold crossover. **(A)** Top-down view of the full edges. The connectivity and nucleotide counts in the structure predictions matches the intended design (schematized in Figure S11 for four helical turns per edge). Because the algorithm generates the structure prediction by affixing the vertex nucleotides for the scaffold on the top and bottom helices and interpolating nucleotide positions between, there is steric clash at crossovers in the output structure drawing. This clash might be avoided by the scaffold nucleotides entering the vertex from a different point in the helical rotation. Blue: scaffold strand. Grey: staple strands. Green: base pairs at crossovers.

Yellow: base pairs at staple nicks. Pink: poly-T loops, bridging to neighboring edges. **(B)** End views of the edges. The atomic model is generated with an assumption that the scaffold (blue) will exit the duplex at a helical position opposite the other duplex of the edge, so that the scaffold is near to its entry point in the subsequent edge and no unpaired scaffold is required.

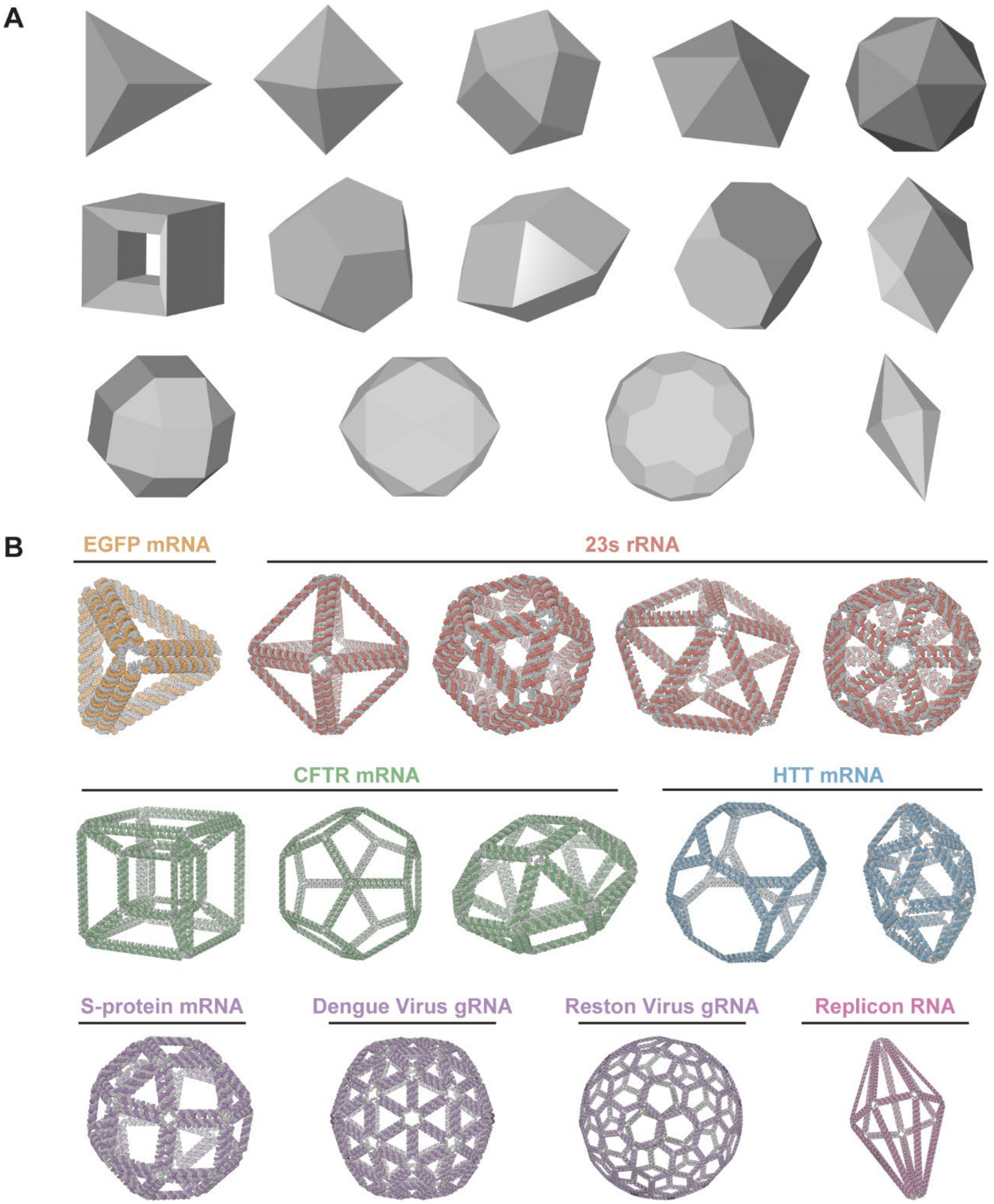

**Figure S25.** 14 A-form objects designed with the pyDAEDALUSX software. **(A)** Input geometries, and **(B)** output 3D structure predictions, using a variety of RNA scaffold sequence inputs. Geometries and edge lengths were chosen for each sequence such that all or a large fragment of the input scaffold sequence would be incorporated into the folded structure design.

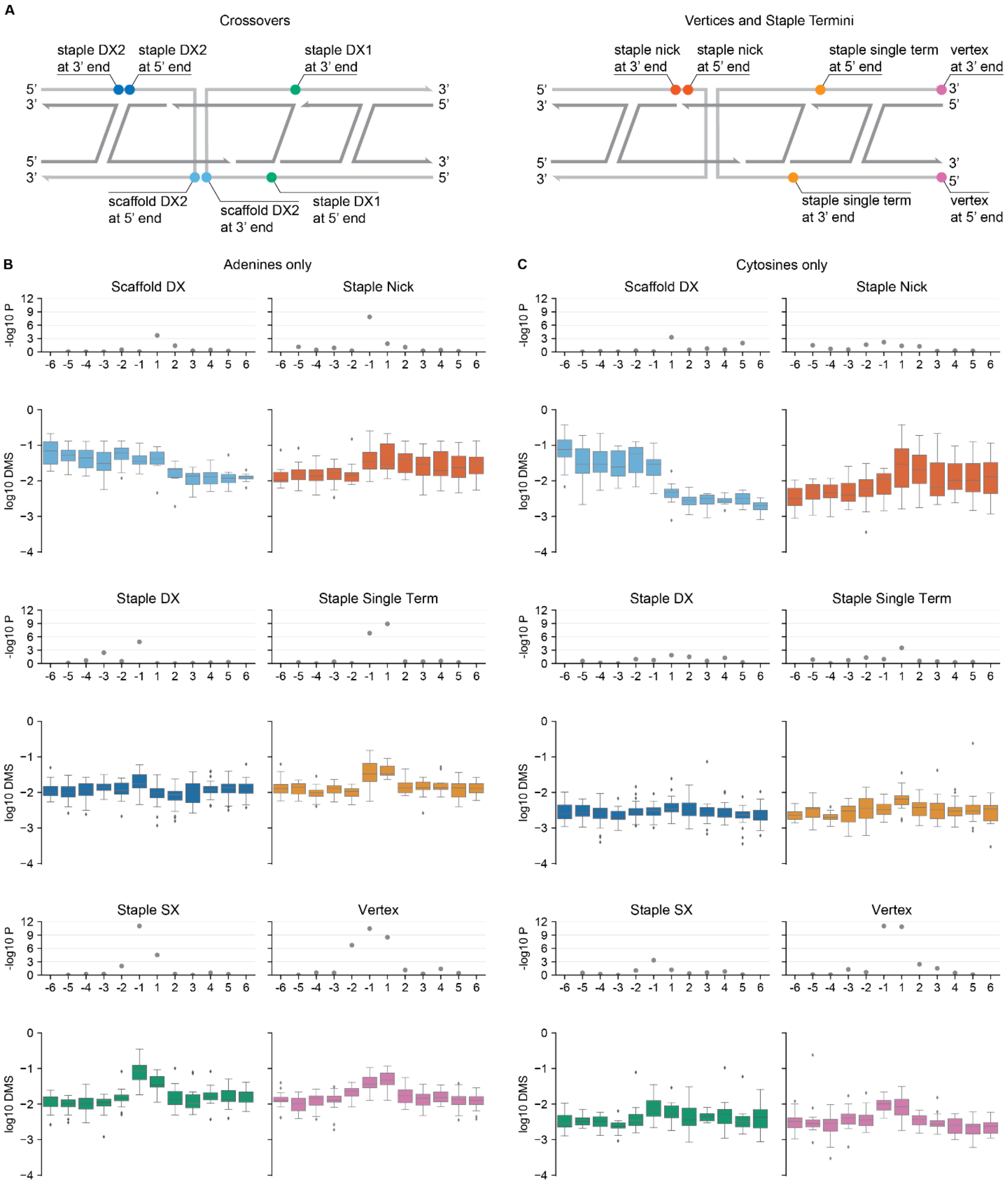

**Figure S26**. DMS reactivities for A-form origami at nucleotides upstream and downstream of features **(A)**, with **(B)** adenines and **(C)** cytosines analyzed separately.

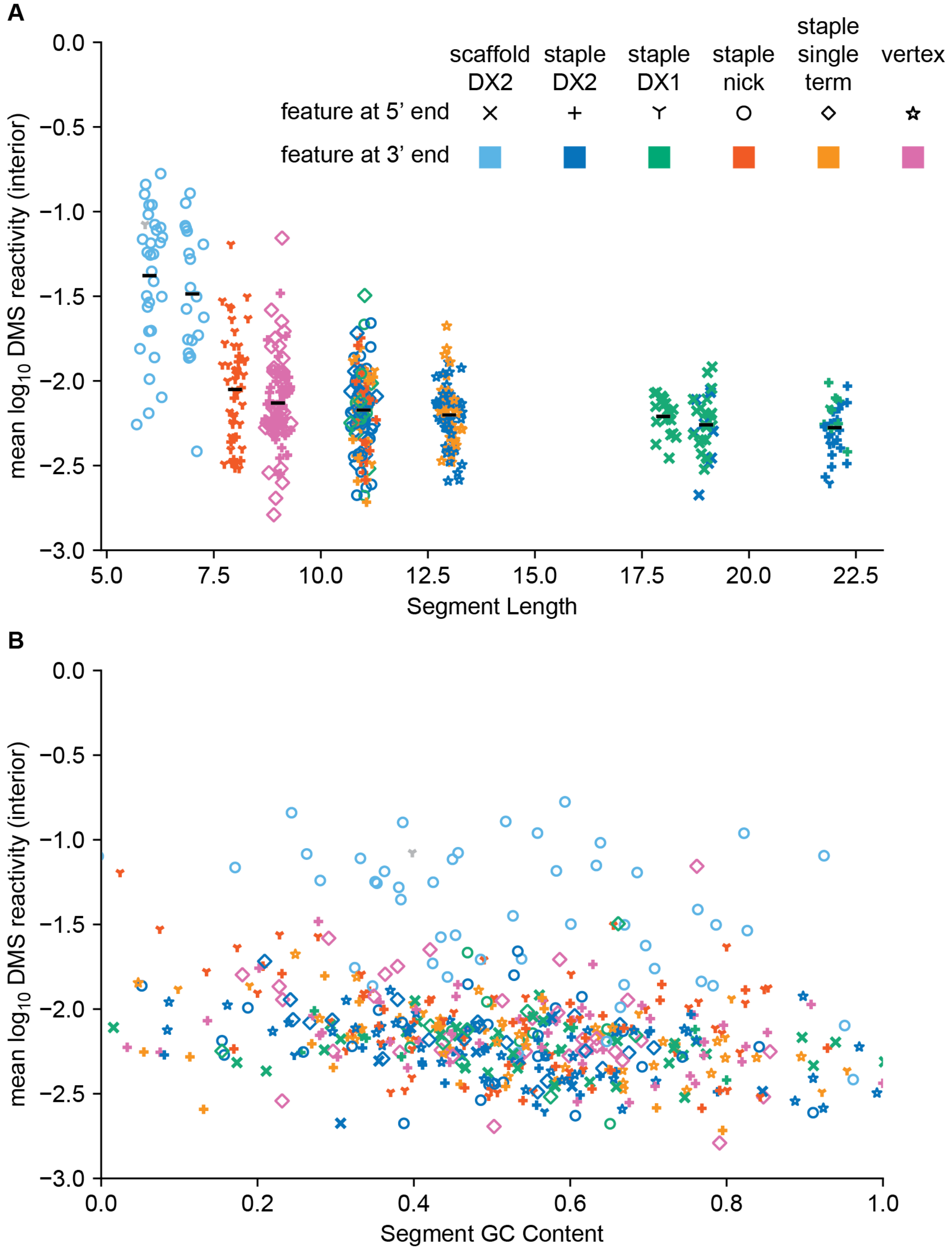

**Figure S27.** Relationship between (A) length or (B) GC content and the mean DMS reactivities over all interior (i.e., excluding the first and last) A and C residues in each A-form origami segment. Each segment corresponds to one point whose color and shape are determined by the structural features at its 3' and 5' ends, respectively.

**Table S1. Template and RNA scaffold sequences.**

| | |
|---|---|
| EGFP gBlock | GCCAGTGAATTCTAATACGACTCACTATAGGTAGCTAAGGAGGTAAATAATGGTGAGCAAGGGCG<br>AGGAGCTGTTCACCGGGGTGGTGCCCATCCTGGTCGAGCTGGACGGCGACGTAAACGGCCACAA<br>GTTCAGCGTGTCCGGCGAGGGCGAGGGCGATGCCACCTACGGCAAGCTGACCCTGAAGTTCATC<br>TGCACCACCGGCAAGCTGCCCGTGCCCTGGCCCACCCTCGTGACCACCCTGACCTACGGCGTGC<br>AGTGCTTCAGCCGCTACCCCGACCACATGAAGCAGCACGACTTCTTCAAGTCCGCCATGCCCGAA<br>GGCTACGTCCAGGAGCGCACCATCTTCTTCAAGGACGACGGCAACTACAAGACCCGCGCCGAGG<br>TGAAGTTCGAGGGCGACACCCTGGTGAACCGCATCGAGCTGAAGGGCATCGACTTCAAGGAGGA<br>CGGCAACATCCTGGGGCACAAGCTGGAGTACAACTACAACAGCCACAACGTCTATATCATGGCCG<br>ACAAGCAGAAGAACGGCATCAAGGTGAACTTCAAGATCCGCCACAACATCGAGGACGGCAGCGT<br>GCAGCTCGCCGACCACTACCAGCAGAACACCCCCATCGGCGACGGCCCCGTGCTGCTGCCCGAC<br>AACCACTACCTGAGCACCCAGTCCGCCCTGAGCAAAGACCCCAACGAGAAGCGCGATCACATGG<br>TCCTGCTGGAGTTCGTGACCGCCGCCGGGATCACTCTCGGCATGGACGAGCTGTACAAGTAACT<br>GCAGGCATGC |
| EGFP-pUC19 | TCGCGCGTTTCGGTGATGACGGTGAAAACCTCTGACACATGCAGCTCCCGGAGACGGTCACAGCT<br>TGTCTGTAAGCGGATGCCGGGAGCAGACAAGCCCGTCAGGGCGCGTCAGCGGGTGTTGGCGGG<br>TGTCGGGGCTGGCTTAACTATGCGGCATCAGAGCAGATTGTACTGAGAGTGCACCATATGCGGTG<br>TGAAATACCGCACAGATGCGTAAGGAGAAAATACCGCATCAGGCGCCATTCGCCATTCAGGCTGC<br>GCAACTGTTGGGAAGGGCGATCGGTGCGGGCCTCTTCGCTATTACGCCAGCTGGCGAAAGGGGG<br>ATGTGCTGCAAGGCGATTAAGTTGGGTAACGCCAGGGTTTTCCCAGTCACGACGTTGTAAAACGA<br>CGGCCAGTGAATTCTAATACGACTCACTATAGGTAGCTAAGGAGGTAAATAATGGTGAGCAAGGG<br>CGAGGAGCTGTTCACCGGGGTGGTGCCCATCCTGGTCGAGCTGGACGGCGACGTAAACGGCCA<br>CAAGTTCAGCGTGTCCGGCGAGGGCGAGGGCGATGCCACCTACGGCAAGCTGACCCTGAAGTTC<br>ATCTGCACCACCGGCAAGCTGCCCGTGCCCTGGCCCACCCTCGTGACCACCCTGACCTACGGCG<br>TGCAGTGCTTCAGCCGCTACCCCGACCACATGAAGCAGCACGACTTCTTCAAGTCCGCCATGCCC<br>GAAGGCTACGTCCAGGAGCGCACCATCTTCTTCAAGGACGACGGCAACTACAAGACCCGCGCCG<br>AGGTGAAGTTCGAGGGCGACACCCTGGTGAACCGCATCGAGCTGAAGGGCATCGACTTCAAGGA<br>GGACGGCAACATCCTGGGGCACAAGCTGGAGTACAACTACAACAGCCACAACGTCTATATCATGG<br>CCGACAAGCAGAAGAACGGCATCAAGGTGAACTTCAAGATCCGCCACAACATCGAGGACGGCAG<br>CGTGCAGCTCGCCGACCACTACCAGCAGAACACCCCCATCGGCGACGGCCCCGTGCTGCTGCCC<br>GACAACCACTACCTGAGCACCCAGTCCGCCCTGAGCAAAGACCCCAACGAGAAGCGCGATCACA<br>TGGTCCTGCTGGAGTTCGTGACCGCCGCCGGGATCACTCTCGGCATGGACGAGCTGTACAAGTA<br>ACTGCAGGCATGCAAGCTTGGCGTAATCATGGTCATAGCTGTTTCCTGTGTGAAATTGTTATCCGC<br>TCACAATTCCACACAACATACGAGCCGGAAGCATAAAGTGTAAAGCCTGGGGTGCCTAATGAGTG<br>AGCTAACTCACATTAATTGCGTTGCGCTCACTGCCCGCTTTCCAGTCGGGAAACCTGTCGTGCCA<br>GCTGCATTAATGAATCGGCCAACGCGCGGGGAGAGGCGGTTTGCGTATTGGGCGCTCTTCCGCT<br>TCCTCGCTCACTGACTCGCTGCGCTCGGTCGTTCGGCTGCGGCGAGCGGTATCAGCTCACTCAAA<br>GGCGGTAATACGGTTATCCACAGAATCAGGGGATAACGCAGGAAAGAACATGTGAGCAAAAGGCC<br>AGCAAAAGGCCAGGAACCGTAAAAAGGCCGCGTTGCTGGCGTTTTTCCATAGGCTCCGCCCCCCT<br>GACGAGCATCACAAAAATCGACGCTCAAGTCAGAGGTGGCGAAACCCGACAGGACTATAAAGATA<br>CCAGGCGTTTCCCCCTGGAAGCTCCCTCGTGCGCTCTCCTGTTCCGACCCTGCCGCTTACCGGAT<br>ACCTGTCCGCCTTTCTCCCTTCGGGAAGCGTGGCGCTTTCTCATAGCTCACGCTGTAGGTATCTCA<br>GTTCGGTGTAGGTCGTTCGCTCCAAGCTGGGCTGTGTGCACGAACCCCCCGTTCAGCCCGACCG<br>CTGCGCCTTATCCGGTAACTATCGTCTTGAGTCCAACCCGGTAAGACACGACTTATCGCCACTGG<br>CAGCAGCCACTGGTAACAGGATTAGCAGAGCGAGGTATGTAGGCGGTGCTACAGAGTTCTTGAAG<br>TGGTGGCCTAACTACGGCTACACTAGAAGAACAGTATTTGGTATCTGCGCTCTGCTGAAGCCAGTT<br>ACCTTCGGAAAAAGAGTTGGTAGCTCTTGATCCGGCAAACAAACCACCGCTGGTAGCGGTGGTTT<br>TTTTGTTTGCAAGCAGCAGATTACGCGCAGAAAAAAAGGATCTCAAGAAGATCCTTTGATCTTTTCT<br>ACGGGGTCTGACGCTCAGTGGAACGAAAACTCACGTTAAGGGATTTTGGTCATGAGATTATCAAAA<br>AGGATCTTCACCTAGATCCTTTTAAATTAAAAATGAAGTTTTAAATCAATCTAAAGTATATATGAGTA<br>AACTTGGTCTGACAGTTACCAATGCTTAATCAGTGAGGCACCTATCTCAGCGATCTGTCTATTTCGT<br>TCATCCATAGTTGCCTGACTCCCCGTCGTGTAGATAACTACGATACGGGAGGGCTTACCATCTGG<br>CCCCAGTGCTGCAATGATACCGCGAGACCCACGCTCACCGGCTCCAGATTTATCAGCAATAAACC<br>AGCCAGCCGGAAGGGCCGAGCGCAGAAGTGGTCCTGCAACTTTATCCGCCTCCATCCAGTCTATT<br>AATTGTTGCCGGGAAGCTAGAGTAAGTAGTTCGCCAGTTAATAGTTTGCGCAACGTTGTTGCCATT<br>GCTACAGGCATCGTGGTGTCACGCTCGTCGTTTGGTATGGCTTCATTCAGCTCCGGTTCCCAACG<br>ATCAAGGCGAGTTACATGATCCCCCATGTTGTGCAAAAAAGCGGTTAGCTCCTTCGGTCCTCCGAT<br>CGTTGTCAGAAGTAAGTTGGCCGCAGTGTTATCACTCATGGTTATGGCAGCACTGCATAATTCTCT<br>TACTGTCATGCCATCCGTAAGATGCTTTTCTGTGACTGGTGAGTACTCAACCAAGTCATTCTGAGA<br>ATAGTGTATGCGGCGACCGAGTTGCTCTTGCCCGGCGTCAATACGGGATAATACCGCGCCACATA<br>GCAGAACTTTAAAAGTGCTCATCATTGGAAAACGTTCTTCGGGGCGAAAACTCTCAAGGATCTTAC |

| | |
|---|---|
| | CGCTGTTGAGATCCAGTTCGATGTAACCCACTCGTGCACCCAACTGATCTTCAGCATCTTTTACTTT<br>CACCAGCGTTTCTGGGTGAGCAAAAACAGGAAGGCAAAATGCCGCAAAAAAGGGAATAAGGGCG<br>ACACGGAAATGTTGAATACTCATACTCTTCCTTTTTCAATATTATTGAAGCATTTATCAGGGTTATTG<br>TCTCATGAGCGGATACATATTTGAATGTATTTAGAAAAATAAACAAATAGGGGTTCCGCGCACATTT<br>CCCCGAAAAGTGCCACCTGACGTCTAAGAAACCATTATTATCATGACATTAACCTATAAAAATAGGC<br>GTATCACGAGGCCCTTTCGTC |
| EGFP RNA scaffold | AAGGGCGAGGAGCUGUUCACCGGGGUGGUGCCCAUCCUGGUCGAGCUGGACGGCGACGUAAA<br>CGGCCACAAGUUCAGCGUGUCCGGCGAGGGCGAGGGCGAUGCCACCUACGGCAAGCUGACCC<br>UGAAGUUCAUCUGCACCACCGGCAAGCUGCCCGUGCCCUGGCCCACCCUCGUGACCACCCUGA<br>CCUACGGCGUGCAGUGCUUCAGCCGCUACCCCGACCACAUGAAGCAGCACGACUUCUUCAAGU<br>CCGCCAUGCCCGAAGGCUACGUCCAGGAGCGCACCAUCUUCUUCAAGGACGACGGCAACUACA<br>AGACCCGCGCCGAGGUGAAGUUCGAGGGCGACACCCUGGUGAACCGCAUCGAGCUGAAGGGC<br>AUCGACUUCAAGGAGGACGGCAACAUCCUGGGGCACAAGCUGGAGUACAACUACAACAGCCAC<br>AACGUCUAUAUCAUGGCCGACAAGCAGAAGAACGGCAUCAAGGUGAACUUCAAGAUCCGCCACA<br>ACAUCGAGGACGGCAGCGUGCAGCUCGCCGACCACUACCAGCAGAACACCCCCAUCGGCGACG<br>GCCCCGUGCUGCUGCCCGACAACCACUACCUGAGCACCCAGUCCGCCCUGAGCAAAGACCCCA<br>ACGAGAAGCGCGAUCACAUGGUCCUGCUGGAGUUCGUGACCGCCGCCGGGAUCACUCUCGGC<br>AUGGACGAGCUGUACAAGUAACUGCAGGCAUGCAAGCUUGGCGUAAUCAUGGUCAUAGCUGUU<br>UCCUGUGUGAAAUUGUUAUCCGCUCACAAUUCCACACAA |
| EGFP RNA sequence | GGUAGCUAAGGAGGUAAAUAAUGGUGAGCAAGGGCGAGGAGCUGUUCACCGGGGUGGUGCCC<br>AUCCUGGUCGAGCUGGACGGCGACGUAAACGGCCACAAGUUCAGCGUGUCCGGCGAGGGCGA<br>GGGCGAUGCCACCUACGGCAAGCUGACCCUGAAGUUCAUCUGCACCACCGGCAAGCUGCCCGU<br>GCCCUGGCCCACCCUCGUGACCACCCUGACCUACGGCGUGCAGUGCUUCAGCCGCUACCCCG<br>ACCACAUGAAGCAGCACGACUUCUUCAAGUCCGCCAUGCCCGAAGGCUACGUCCAGGAGCGCA<br>CCAUCUUCUUCAAGGACGACGGCAACUACAAGACCCGCGCCGAGGUGAAGUUCGAGGGCGACA<br>CCCUGGUGAACCGCAUCGAGCUGAAGGGCAUCGACUUCAAGGAGGACGGCAACAUCCUGGGG<br>CACAAGCUGGAGUACAACUACAACAGCCACAACGUCUAUAUCAUGGCCGACAAGCAGAAGAACG<br>GCAUCAAGGUGAACUUCAAGAUCCGCCACAACAUCGAGGACGGCAGCGUGCAGCUCGCCGACC<br>ACUACCAGCAGAACACCCCCAUCGGCGACGGCCCCGUGCUGCUGCCCGACAACCACUACCUGA<br>GCACCCAGUCCGCCCUGAGCAAAGACCCCAACGAGAAGCGCGAUCACAUGGUCCUGCUGGAGU<br>UCGUGACCGCCGCCGGGAUCACUCUCGGCAUGGACGAGCUGUACAAGUAACUGCAGGCAUGCA<br>AGCUUGGCGUAAUCAUGGUCAUAGCUGUUUCCUGUGUGAAAUUGUUAUCCGCUCACAAUUCCA<br>CACAACAUACG |
| EGFP DNA template | GAATTCTAATACGACTCACTATAGGTAGCTAAGGAGGTAAATAATGGTGAGCAAGGGCGAGGAGC<br>TGTTCACCGGGGTGGTGCCCATCCTGGTCGAGCTGGACGGCGACGTAAACGGCCACAAGTTCAG<br>CGTGTCCGGCGAGGGCGAGGGCGATGCCACCTACGGCAAGCTGACCCTGAAGTTCATCTGCACC<br>ACCGGCAAGCTGCCCGTGCCCTGGCCCACCCTCGTGACCACCCTGACCTACGGCGTGCAGTGCT<br>TCAGCCGCTACCCCGACCACATGAAGCAGCACGACTTCTTCAAGTCCGCCATGCCCGAAGGCTAC<br>GTCCAGGAGCGCACCATCTTCTTCAAGGACGACGGCAACTACAAGACCCGCGCCGAGGTGAAGT<br>TCGAGGGCGACACCCTGGTGAACCGCATCGAGCTGAAGGGCATCGACTTCAAGGAGGACGGCAA<br>CATCCTGGGGCACAAGCTGGAGTACAACTACAACAGCCACAACGTCTATATCATGGCCGACAAGC<br>AGAAGAACGGCATCAAGGTGAACTTCAAGATCCGCCACAACATCGAGGACGGCAGCGTGCAGCT<br>CGCCGACCACTACCAGCAGAACACCCCCATCGGCGACGGCCCCGTGCTGCTGCCCGACAACCAC<br>TACCTGAGCACCCAGTCCGCCCTGAGCAAAGACCCCAACGAGAAGCGCGATCACATGGTCCTGC<br>TGGAGTTCGTGACCGCCGCCGGGATCACTCTCGGCATGGACGAGCTGTACAAGTAACTGCAGGC<br>ATGCAAGCTTGGCGTAATCATGGTCATAGCTGTTTCCTGTGTGAAATTGTTATCCGCTCACAATTCC<br>ACACAACATACG |
| rrlB RNA scaffold input sequence (23S) | GCCCTGGCAGTCAGAGGCGATGAAGGACGTGCTAATCTGCGATAAGCGTCGGTAAGGTGATATG<br>AACCGTTATAACCGGCGATTTCCGAATGGGGAAACCCAGTGTGTTTCGACACACTATCATTAACTG<br>AATCCATAGGTTAATGAGGCGAACCGGGGGAACTGAAACATCTAAGTACCCCGAGGAAAAGAAAT<br>CAACCGAGATTCCCCCAGTAGCGGCGAGCGAACGGGGAGCAGCCCAGAGCCTGAATCAGTGTGT<br>GTGTTAGTGGAAGCGTCTGGAAAGGCGCGCGATACAGGGTGACAGCCCCGTACACAAAAATGCA<br>CATGCTGTGAGCTCGATGAGTAGGGCGGGACACGTGGTATCCTGTCTGAATATGGGGGGACCAT<br>CCTCCAAGGCTAAATACTCCTGACTGACCGATAGTGAACCAGTACCGTGAGGGAAAGGCGAAAAG<br>AACCCCGGCGAGGGGAGTGAAAAAGAACCTGAAACCGTGTACGTACAAGCAGTGGGAGCACGCT<br>TAGGCGTGTGACTGCGTACCTTTTGTATAATGGGTCAGCGACTTATATTCTGTAGCAAGGTTAACC<br>GAATAGGGGAGCCGAAGGGAAACCGAGTCTTAACTGGGCGTTAAGTTGCAGGGTATAGACCCGA<br>AACCCGGTGATCTAGCCATGGGCAGGTTGAAGGTTGGGTAACACTAACTGGAGGACCGAACCGA<br>CTAATGTTGAAAAATTAGCGGATGACTTGTGGCTGGGGGTGAAAGGCCAATCAAACCGGGAGATA<br>GCTGGTTCTCCCCGAAAGCTATTTAGGTAGCGCCTCGTGAATTCATCTCCGGGGGTAGAGCACTG |

| | |
|---|---|
| | TTTCGGCAAGGGGGTCATCCCGACTTACCAACCCGATGCAAACTGCGAATACCGGAGAATGTTAT<br>CACGGGAGACACACGGCGGGTGCTAACGTCCGTCGTGAAGAGGGAAACAACCCAGACCGCCAG<br>CTAAGGTCCCAAAGTCATGGTTAAGTGGGAAACGATGTGGGAAGGCCCAGACAGCCAGGATGTTG<br>GCTTAGAAGCAGCCATCATTTAAAGAAAGCGTAATAGCTCACTGGTCGAGTCGGCCTGCGCGGAA<br>GATGTAACGGGGCTAAACCATGCACCGAAGCTGCGGCAGCGACGCTTATGCGTTGTTGGGTAGG<br>GGAGCGTTCTGTAAGCCTGCGAAGGTGTGCTGTGAGGCATGCTGGAGGTATCAGAAGTGCGAAT<br>GCTGACATAAGTAACGATAAAGCGGGTGAAAAGCCCGCTCGCCGGAAGACCAAGGGTTCCTGTC<br>CAACGTTAATCGGGGCAGGGTGAGTCGACCCCTAAGGCGAGGCCGAAAGGCGTAGTCGATGGGA<br>AACAGGTTAATATTCCTGTACTTGGTGTTACTGCGAAGGGGGGACGGAGAAGGCTATGTTGGCCG<br>GGCGACGGTTGTCCCGGTTTAAGCGTGTAGGCTGGTTTTCCAGGCAAATCCGGAAAATCAAGGCT<br>GAGGCGTGATGACGAGGCACTACGGTGCTGAAGCAACAAATGCCCTGCTTCCAGGAAAAGCCTCT<br>AAGCATCAGGTAACATCAAATCGTACCCCAAACCGACACAGGTGGTCAGGTAGAGAATACCAAGG<br>CGCTTGAGAGAACTCGGGTGAAGGAACTAGGCAAAATGGTGCCGTAACTTCGGGAGAAGGCACG<br>CTGATATGTAGGTGAAGCGACTTGCTCGTGGAGCTGAAATCAGTCGAAGATACCAGCTGGCTGCA<br>ACTGTTTATTAAAAACACAGCACTGTGCAAACACGAAAGTGGACGTATACGGTGTGACGCCTGCCC<br>GGTGCCGGAAGGTTAATTGATGGGGTTAGCGCAAGCGAAGCTCTTGATCGAAGCCCCGGTAAAC<br>GGCGGCCGTAACTATAACGGTCCTAAGGTAGCGAAATTCCTTGTCGGGTAAGTTCCGACCTGCAC<br>GAATGGCGTAATGATGGCCAGGCTGTCTCCACCCGAGACTCA |
| rrlB RNA sequence (23S) | GCCCTGGCAGTCAGAGGCGATGAAGGACGTGCTAATCTGCGATAAGCGTCGGTAAGGTGATATG<br>AACCGTTATAACCGGCGATTTCCGAATGGGGAAACCCAGTGTGTTTCGACACACTATCATTAACTG<br>AATCCATAGGTTAATGAGGCGAACCGGGGGAACTGAAACATCTAAGTACCCCGAGGAAAAGAAAT<br>CAACCGAGATTCCCCCAGTAGCGGCGAGCGAACGGGGAGCAGCCCAGAGCCTGAATCAGTGTGT<br>GTGTTAGTGGAAGCGTCTGGAAAGGCGCGCGATACAGGGTGACAGCCCCGTACACAAAAATGCA<br>CATGCTGTGAGCTCGATGAGTAGGGCGGGACACGTGGTATCCTGTCTGAATATGGGGGGACCAT<br>CCTCCAAGGCTAAATACTCCTGACTGACCGATAGTGAACCAGTACCGTGAGGGAAAGGCGAAAAG<br>AACCCCGGCGAGGGGAGTGAAAAAGAACCTGAAACCGTGTACGTACAAGCAGTGGGAGCACGCT<br>TAGGCGTGTGACTGCGTACCTTTTGTATAATGGGTCAGCGACTTATATTCTGTAGCAAGGTTAACC<br>GAATAGGGGAGCCGAAGGGAAACCGAGTCTTAACTGGGCGTTAAGTTGCAGGGTATAGACCCGA<br>AACCCGGTGATCTAGCCATGGGCAGGTTGAAGGTTGGGTAACACTAACTGGAGGACCGAACCGA<br>CTAATGTTGAAAAATTAGCGGATGACTTGTGGCTGGGGGTGAAAGGCCAATCAAACCGGGAGATA<br>GCTGGTTCTCCCCGAAAGCTATTTAGGTAGCGCCTCGTGAATTCATCTCCGGGGGTAGAGCACTG<br>TTTCGGCAAGGGGGTCATCCCGACTTACCAACCCGATGCAAACTGCGAATACCGGAGAATGTTAT<br>CACGGGAGACACACGGCGGGTGCTAACGTCCGTCGTGAAGAGGGAAACAACCCAGACCGCCAG<br>CTAAGGTCCCAAAGTCATGGTTAAGTGGGAAACGATGTGGGAAGGCCCAGACAGCCAGGATGTTG<br>GCTTAGAAGCAGCCATCATTTAAAGAAAGCGTAATAGCTCACTGGTCGAGTCGGCCTGCGCGGAA<br>GATGTAACGGGGCTAAACCATGCACCGAAGCTGCGGCAGCGACGCTTATGCGTTGTTGGGTAGG<br>GGAGCGTTCTGTAAGCCTGCGAAGGTGTGCTGTGAGGCATGCTGGAGGTATCAGAAGTGCGAAT<br>GCTGACATAAGTAACGATAAAGCGGGTGAAAAGCCCGCTCGCCGGAAGACCAAGGGTTCCTGTC<br>CAACGTTAATCGGGGCAGGGTGAGTCGACCCCTAAGGCGAGGCCGAAAGGCGTAGTCGATGGGA<br>AACAGGTTAATATTCCTGTACTTGGTGTTACTGCGAAGGGGGGACGGAGAAGGCTATGTTGGCCG<br>GGCGACGGTTGTCCCGGTTTAAGCGTGTAGGCTGGTTTTCCAGGCAAATCCGGAAAATCAAGGCT<br>GAGGCGTGATGACGAGGCACTACGGTGCTGAAGCAACAAATGCCCTGCTTCCAGGAAAAGCCTCT<br>AAGCATCAGGTAACATCAAATCGTACCCCAAACCGACACAGGTGGTCAGGTAGAGAATACCAAGG<br>CGCTTGAGAGAACTCGGGTGAAGGAACTAGGCAAAATGGTGCCGTAACTTCGGGAGAAGGCACG<br>CTGATATGTAGGTGAGGTCCCTCGCGGATGGAGCTGAAATCAGTCGAAGATACCAGCTGGCTGCA<br>ACTGTTTATTAAAAACACAGCACTGTGCAAACACGAAAGTGGACGTATACGGTGTGACGCCTGCCC<br>GGTGCCGGAAGGTTAATTGATGGGGTTAGCGCAAGCGAAGCTCTTGATCGAAGCCCCGGTAAAC<br>GGCGGCCGTAACTATAACGGTCCTAAGGTAGCGAAATTCCTTGTCGGGTAAGTTCCGACCTGCAC<br>GAATGGCGTAATGATGGCCAGGCTGTCTCCACCCGAGACTCA |
| rrlB DNA template | pCW1 plasmid from:<br>Carl J. Weitzmann, Philip R. Cunningham, James Ofengand, Cloning, in vitro transcription, and biological activity of Escherichia coli 23S ribosomal RNA, Nucleic Acids Research, Volume 18, Issue 12, 25 June 1990, Pages 3515–3520, https://doi.org/10.1093/nar/18.12.3515 |
| M13 DNA source | M13mp18 ssDNA from New England Biolabs |
| M13 O44 RNA scaffold | GUCUCACUGGUGAAAAGAAAAACCACCCUGGCGCCCAAUACGCAAACCGCCUCUCCCCGCGCG<br>UUGGCCGAUUCAUUAAUGCAGCUGGCACGACAGGUUUCCCGACUGGAAAGCGGGCAGUGAGC<br>GCAACGCAAUUAAUGUGAGUUAGCUCACUCAUUAGGCACCCCAGGCUUUACACUUUAUGCUUC<br>CGGCUCGUAUGUUGUGUGGAAUUGUGAGCGGAUAACAAUUUCACACAGGAAACAGCUAUGACC |

| | |
|---|---|
| | AUGAUUACGAAUUCGAGCUCGGUACCCGGGGAUCCUCUAGAGUCGACCUGCAGGCAUGCAAGC<br>UUGGCACUGGCCGUCGUUUUACAACGUCGUGACUGGGAAAACCCUGGCGUUACCCAACUUAAU<br>CGCCUUGCAGCACAUCCCCCUUUCGCCAGCUGGCGUAAUAGCGAAGAGGCCCGCACCGAUCGC<br>CCUUCCCAACAGUUGCGCAGCCUGAAUGGCGAAUGGCGCUUUGCCUGGUUUCCGGCACCAGAA<br>GCGGUGCCGGAAAGCUGGCUGGAGUGCGAUCUUCCUGAGGCCGAUACUGUCGUCGUCCCCUC<br>AAACUGGCAGAUGCACGGUUACGAUGCGCCCAUCUACACCAACGUGACCUAUCCCAUUACGGU<br>CAAUCCGCCGUUUGUUCCCACGGAGAAUCCGACGGGUUGUUACUCGCUCACAUUUAAUGUUGA<br>UGAAAGCUGGCUACAGGAAGGCCAGACGCGAAUUAUUUUUGAUGGCGUUCCUAUUGGUUAAAA<br>AAUGAGCUGAUUUAACAAAAAUUUAAUGCGAAUUUUAACAAAAUAUUAACGUUUACAAUUUAAAU<br>AUUUGCUUAUACAAUCUUCCUGUUUUUGGGGCUUUUCUGAUUAUCAACCGGGGUACAUAUGAU<br>UGACAUGCUAGUUUUACGAUUACCGUUCAUCGAUUCUCUUGUUUGCUCCAGACUCUCAGGCAA<br>UGACCUGAUAGCCUUUGUAGAUCUCUCAAAAAUAGCUACCCUCUCCGGCAUUAAUUUAUCAGCU<br>AGAACGGUUGAAUAUCAUAUUGAUGGUGAUUUGACUGUCUCCGGCCU |
| M13<br>O44<br>RNA<br>sequence | GGGUCUCACUGGUGAAAAGAAAAACCACCCUGGCGCCCAAUACGCAAACCGCCUCUCCCCGCG<br>CGUUGGCCGAUUCAUUAAUGCAGCUGGCACGACAGGUUUCCCGACUGGAAAGCGGGCAGUGA<br>GCGCAACGCAAUUAAUGUGAGUUAGCUCACUCAUUAGGCACCCCAGGCUUUACACUUUAUGCU<br>UCCGGCUCGUAUGUUGUGUGGAAUUGUGAGCGGAUAACAAUUUCACACAGGAAACAGCUAUGA<br>CCAUGAUUACGAAUUCGAGCUCGGUACCCGGGGAUCCUCUAGAGUCGACCUGCAGGCAUGCAA<br>GCUUGGCACUGGCCGUCGUUUUACAACGUCGUGACUGGGAAAACCCUGGCGUUACCCAACUUA<br>AUCGCCUUGCAGCACAUCCCCCUUUCGCCAGCUGGCGUAAUAGCGAAGAGGCCCGCACCGAUC<br>GCCCUUCCCAACAGUUGCGCAGCCUGAAUGGCGAAUGGCGCUUUGCCUGGUUUCCGGCACCA<br>GAAGCGGUGCCGGAAAGCUGGCUGGAGUGCGAUCUUCCUGAGGCCGAUACUGUCGUCGUCCC<br>CUCAAACUGGCAGAUGCACGGUUACGAUGCGCCCAUCUACACCAACGUGACCUAUCCCAUUAC<br>GGUCAAUCCGCCGUUUGUUCCCACGGAGAAUCCGACGGGUUGUUACUCGCUCACAUUUAAUGU<br>UGAUGAAAGCUGGCUACAGGAAGGCCAGACGCGAAUUAUUUUUGAUGGCGUUCCUAUUGGUUA<br>AAAAAUGAGCUGAUUUAACAAAAAUUUAAUGCGAAUUUUAACAAAAUAUUAACGUUUACAAUUUA<br>AAUAUUUGCUUAUACAAUCUUCCUGUUUUUGGGGCUUUUCUGAUUAUCAACCGGGGUACAUAU<br>GAUUGACAUGCUAGUUUUACGAUUACCGUUCAUCGAUUCUCUUGUUUGCUCCAGACUCUCAGG<br>CAAUGACCUGAUAGCCUUUGUAGAUCUCUCAAAAAUAGCUACCCUCUCCGGCAUUAAUUUAUCA<br>GCUAGAACGGUUGAAUAUCAUAUUGAUGGUGAUUUGACUGUCUCCGGCC |
| M13<br>O44<br>DNA<br>template | GAATTCTAATACGACTCACTATAGGGTCTCACTGGTGAAAAGAAAAACCACCCTGGCGCCCAATAC<br>GCAAACCGCCTCTCCCCGCGCGTTGGCCGATTCATTAATGCAGCTGGCACGACAGGTTTCCCGAC<br>TGGAAAGCGGGCAGTGAGCGCAACGCAATTAATGTGAGTTAGCTCACTCATTAGGCACCCCAGGC<br>TTTACACTTTATGCTTCCGGCTCGTATGTTGTGTGGAATTGTGAGCGGATAACAATTTCACACAGG<br>AAACAGCTATGACCATGATTACGAATTCGAGCTCGGTACCCGGGGATCCTCTAGAGTCGACCTGC<br>AGGCATGCAAGCTTGGCACTGGCCGTCGTTTTACAACGTCGTGACTGGGAAAACCCTGGCGTTAC<br>CCAACTTAATCGCCTTGCAGCACATCCCCCTTTCGCCAGCTGGCGTAATAGCGAAGAGGCCCGCA<br>CCGATCGCCCTTCCCAACAGTTGCGCAGCCTGAATGGCGAATGGCGCTTTGCCTGGTTTCCGGCA<br>CCAGAAGCGGTGCCGGAAAGCTGGCTGGAGTGCGATCTTCCTGAGGCCGATACTGTCGTCGTCC<br>CCTCAAACTGGCAGATGCACGGTTACGATGCGCCCATCTACACCAACGTGACCTATCCCATTACG<br>GTCAATCCGCCGTTTGTTCCCACGGAGAATCCGACGGGTTGTTACTCGCTCACATTTAATGTTGAT<br>GAAAGCTGGCTACAGGAAGGCCAGACGCGAATTATTTTTGATGGCGTTCCTATTGGTTAAAAAATG<br>AGCTGATTTAACAAAAATTTAATGCGAATTTTAACAAAATATTAACGTTTACAATTTAAATATTTGCTT<br>ATACAATCTTCCTGTTTTTGGGGCTTTTCTGATTATCAACCGGGGTACATATGATTGACATGCTAGT<br>TTTACGATTACCGTTCATCGATTCTCTTGTTTGCTCCAGACTCTCAGGCAATGACCTGATAGCCTTT<br>GTAGATCTCTCAAAAATAGCTACCCTCTCCGGCATTAATTTATCAGCTAGAACGGTTGAATATCATA<br>TTGATGGTGATTTGACTGTCTCCGGCC |
| rsc1218<br>v1<br>gblock | GGGCCTAGTGAGTGCATTAGAGACTCAAGCCATGTATCCATGACCAGAAGAGGGGACCCTAGGC<br>CAAGGACATGAGGGGCTCTATGCACTAATGTCTAGTCAGGCCTTCACCCTAGTCATCTCTATGTGG<br>AATACAATTGGGATTCATGAGAGAATCATGCAAGCCAAAGTCAGCAGGTAGCCTTATTTGATGGAG<br>TTAACCAACAGCCAAATTTATTGCCCAGAGCCCATTAACCATGCTAGAGGAGGGTTTCTCCTGATG<br>ACTGGCTCCCTGACATCTGTGTCTGGGCCTCAAATTGGTTTGAAGCTGATGGGCTATGCATAATCC<br>CATCACTAAGACCTTGGTCTTGCTTGCTAGAACTTTTCATTACACCACTAGCTGAAGTTTGTTATGA<br>CTGCCCCTAATGGCAGGCTCCTCAGGGACAGAAGGAACTGGCCAACCCCATATCTAATGTTGTTG<br>GGCTTGCTGTAGGGTACCCATCTATGAGGAGGCTTGTAACTGTGATGCTCCCTTAGCTACTTTTGA<br>TGTATATGCACAGTGTTGGCAGTGGACCTTGCTCCATGCAATTCATATGGCTATCTTGCAGTCCAT<br>GGCCCACCCCAGACTTCATTGCTGCCCTGTGACACCATTCATTAGAACCTGGTATGGAAGTGTAC<br>CACAGAGCTGCAAGGGGTTGTAGACCACTTATGGTGAAATTTGTAGAATCCAGGGTGGTGAGCTG<br>TAAATGAAACTTTGGATAACTGGCTTTCTAGAGCCTTCCTCCCACCCTCCCCTGTTTGCCATGCCC<br>CAGGGGCAAGCACTGCCCTTTGCACTCTCACTACAGGTAAAGAGTGTCCCCAGGAGTAAATTTCT |

| | |
|---|---|
| | AGCCCCATAGAAAAGGAAGGTCTAGAGGGAAATTGGCAATGGGCACCTGTCCCATTATAAGCATC<br>TATTTGAAGATGCCTGGCACAATAGATCAGTAGGGGAGTTGCTATACCCACTGGTAAGGTTAGATG<br>GTGTTAGTAGGCAGGTCAGCACAGACCTGCACATACAGCATGGGTGGTTACCCAACCATACACAA<br>GACCCATGAAAGTGTGGAAGCTCTTGTACACCCTCTGGCACCAGCATAGTGTTGCATTGTGCACT<br>GAAAGAGCTACCCCAGGCTGTAGCCTAATTTCTTATGGGGATAGGGGCCTTGCAATTTCAAAGCCT<br>TGGACACCTGACCCCACCTCTGAGGGTCTCAGGCCCTC |
| rsc1218 v1_rT55 RNA scaffold | GGGCCUAGUGAGUGCAUUAGAGACUCAAGCCAUGUAUCCAUGACCAGAAGAGGGGACCCUAGG<br>CCAAGGACAUGAGGGGCUCUAUGCACUAAUGUCUAGUCAGGCCUUCACCCUAGUCAUCUCUAU<br>GUGGAAUACAAUUGGGAUUCAUGAGAGAAUCAUGCAAGCCAAAGUCAGCAGGUAGCCUUAUUU<br>GAUGGAGUUAACCAACAGCCAAAUUUAUUGCCCAGAGCCCAUUAACCAUGCUAGAGGAGGGUU<br>UCUCCUGAUGACUGGCUCCCUGACAUCUGUGUCUGGGCCUCAAAUUGGUUUGAAGCUGAUGG<br>GCUAUGCAUAAUCCCAUCACUAAGACCUUGGUCUUGCUUGCUAGAACUUUUCAUUACACCACUA<br>GCUGAAGUUUGUUAUGACUGCCCCUAAUGGCAGGCUCCUCAGGGACAGAAGGAACUGGCCAAC<br>CCCAUAUCUAAUGUUGUUGGGCUUGCUGUAGGGUACCCAUCUAUGAGGAGGCUUGUAACUGU<br>GAUGCUCCCUUAGCUACUUUUGAUGUAUAUGCACAGUGUUGGCAGUGGACCUUGCUCCAUGCA<br>AUUCAUAUGGCUAUCUUGCAGUCCAUGGCCCACCCCAGACUUCAUUGCUGCCCUGUGACACCA<br>UUCAUUAGAACCUGGUAUGGAAGUGUACCAC |
| rsc1218 v1_rT55 RNA sequence | GGGCCUAGUGAGUGCAUUAGAGACUCAAGCCAUGUAUCCAUGACCAGAAGAGGGGACCCUAGG<br>CCAAGGACAUGAGGGGCUCUAUGCACUAAUGUCUAGUCAGGCCUUCACCCUAGUCAUCUCUAU<br>GUGGAAUACAAUUGGGAUUCAUGAGAGAAUCAUGCAAGCCAAAGUCAGCAGGUAGCCUUAUUU<br>GAUGGAGUUAACCAACAGCCAAAUUUAUUGCCCAGAGCCCAUUAACCAUGCUAGAGGAGGGUU<br>UCUCCUGAUGACUGGCUCCCUGACAUCUGUGUCUGGGCCUCAAAUUGGUUUGAAGCUGAUGG<br>GCUAUGCAUAAUCCCAUCACUAAGACCUUGGUCUUGCUUGCUAGAACUUUUCAUUACACCACUA<br>GCUGAAGUUUGUUAUGACUGCCCCUAAUGGCAGGCUCCUCAGGGACAGAAGGAACUGGCCAAC<br>CCCAUAUCUAAUGUUGUUGGGCUUGCUGUAGGGUACCCAUCUAUGAGGAGGCUUGUAACUGU<br>GAUGCUCCCUUAGCUACUUUUGAUGUAUAUGCACAGUGUUGGCAGUGGACCUUGCUCCAUGCA<br>AUUCAUAUGGCUAUCUUGCAGUCCAUGGCCCACCCCAGACUUCAUUGCUGCCCUGUGACACCA<br>UUCAUUAGAACCUGGUAUGGAAGUGUACCAC |
| rsc1218 v1_rT55 DNA template | GGCTTATCGAAATTAATACGACTCACTATAGGGCCTAGTGAGTGCATTAGAGACTCAAGCCATGTA<br>TCCATGACCAGAAGAGGGGACCCTAGGCCAAGGACATGAGGGGCTCTATGCACTAATGTCTAGTC<br>AGGCCTTCACCCTAGTCATCTCTATGTGGAATACAATTGGGATTCATGAGAGAATCATGCAAGCCA<br>AAGTCAGCAGGTAGCCTTATTTGATGGAGTTAACCAACAGCCAAATTTATTGCCCAGAGCCCATTA<br>ACCATGCTAGAGGAGGGTTTCTCCTGATGACTGGCTCCCTGACATCTGTGTCTGGGCCTCAAATT<br>GGTTTGAAGCTGATGGGCTATGCATAATCCCATCACTAAGACCTTGGTCTTGCTTGCTAGAACTTT<br>TCATTACACCACTAGCTGAAGTTTGTTATGACTGCCCCTAATGGCAGGCTCCTCAGGGACAGAAG<br>GAACTGGCCAACCCCATATCTAATGTTGTTGGGCTTGCTGTAGGGTACCCATCTATGAGGAGGCTT<br>GTAACTGTGATGCTCCCTTAGCTACTTTTGATGTATATGCACAGTGTTGGCAGTGGACCTTGCTCC<br>ATGCAATTCATATGGCTATCTTGCAGTCCATGGCCCACCCCAGACTTCATTGCTGCCCTGTGACAC<br>CATTCATTAGAACCTGGTATGGAAGTGTACCAC |
| rsc1218 v1_rT77 RNA scaffold | GGGCCUAGUGAGUGCAUUAGAGACUCAAGCCAUGUAUCCAUGACCAGAAGAGGGGACCCUAGG<br>CCAAGGACAUGAGGGGCUCUAUGCACUAAUGUCUAGUCAGGCCUUCACCCUAGUCAUCUCUAU<br>GUGGAAUACAAUUGGGAUUCAUGAGAGAAUCAUGCAAGCCAAAGUCAGCAGGUAGCCUUAUUU<br>GAUGGAGUUAACCAACAGCCAAAUUUAUUGCCCAGAGCCCAUUAACCAUGCUAGAGGAGGGUU<br>UCUCCUGAUGACUGGCUCCCUGACAUCUGUGUCUGGGCCUCAAAUUGGUUUGAAGCUGAUGG<br>GCUAUGCAUAAUCCCAUCACUAAGACCUUGGUCUUGCUUGCUAGAACUUUUCAUUACACCACUA<br>GCUGAAGUUUGUUAUGACUGCCCCUAAUGGCAGGCUCCUCAGGGACAGAAGGAACUGGCCAAC<br>CCCAUAUCUAAUGUUGUUGGGCUUGCUGUAGGGUACCCAUCUAUGAGGAGGCUUGUAACUGU<br>GAUGCUCCCUUAGCUACUUUUGAUGUAUAUGCACAGUGUUGGCAGUGGACCUUGCUCCAUGCA<br>AUUCAUAUGGCUAUCUUGCAGUCCAUGGCCCACCCCAGACUUCAUUGCUGCCCUGUGACACCA<br>UUCAUUAGAACCUGGUAUGGAAGUGUACCACAGAGCUGCAAGGGGUUGUAGACCACUUAUGGU<br>GAAAUUUGUAGAAUCCAGGGUGGUGAGCUGUAAAUGAAACUUUGGAUAACUGGCUUUCUAGAG<br>CCUUCCUCCCACCCUCCCCUGUUUGCCAUGCCCCAGGGGCAAGCACUGCCCUUUGCACUCUCA<br>CUACAGGUAAAGAGUGUCCCCAGGAGUAAAUUUCUAGCCCCAUAGAAAAGGAAGGUCUAGAGG<br>GAAAUUGGCAAUGGGCACCUGUCCCAUUAUAAGCAUCUAUUUG |
| rsc1218 v1_rT77 RNA sequence | GGGCCUAGUGAGUGCAUUAGAGACUCAAGCCAUGUAUCCAUGACCAGAAGAGGGGACCCUAGG<br>CCAAGGACAUGAGGGGCUCUAUGCACUAAUGUCUAGUCAGGCCUUCACCCUAGUCAUCUCUAU<br>GUGGAAUACAAUUGGGAUUCAUGAGAGAAUCAUGCAAGCCAAAGUCAGCAGGUAGCCUUAUUU<br>GAUGGAGUUAACCAACAGCCAAAUUUAUUGCCCAGAGCCCAUUAACCAUGCUAGAGGAGGGUU<br>UCUCCUGAUGACUGGCUCCCUGACAUCUGUGUCUGGGCCUCAAAUUGGUUUGAAGCUGAUGG<br>GCUAUGCAUAAUCCCAUCACUAAGACCUUGGUCUUGCUUGCUAGAACUUUUCAUUACACCACUA |

| | |
|---|---|
| | GCUGAAGUUUGUUAUGACUGCCCCUAAUGGCAGGCUCCUCAGGGACAGAAGGAACUGGCCAAC<br>CCCAUAUCUAAUGUUGUUGGGCUUGCUGUAGGGUACCCAUCUAUGAGGAGGCUUGUAACUGU<br>GAUGCUCCCUUAGCUACUUUUGAUGUAUAUGCACAGUGUUGGCAGUGGACCUUGCUCCAUGCA<br>AUUCAUAUGGCUAUCUUGCAGUCCAUGGCCCACCCCAGACUUCAUUGCUGCCCUGUGACACCA<br>UUCAUUAGAACCUGGUAUGGAAGUGUACCACAGAGCUGCAAGGGGUUGUAGACCACUUAUGGU<br>GAAAUUUGUAGAAUCCAGGGUGGUGAGCUGUAAAUGAAACUUUGGAUAACUGGCUUUCUAGAG<br>CCUUCCUCCCACCCUCCCCUGUUUGCCAUGCCCCAGGGGCAAGCACUGCCCUUUGCACUCUCA<br>CUACAGGUAAAGAGUGUCCCCAGGAGUAAAUUUCUAGCCCCAUAGAAAAGGAAGGUCUAGAGG<br>GAAAUUGGCAAUGGGCACCUGUCCCAUUAUAAGCAUCUAUUUG |
| rsc1218<br>v1_rT77<br>DNA<br>templat<br>e | GGCTTATCGAAATTAATACGACTCACTATAGGGCCTAGTGAGTGCATTAGAGACTCAAGCCATGTA<br>TCCATGACCAGAAGAGGGGACCCTAGGCCAAGGACATGAGGGGCTCTATGCACTAATGTCTAGTC<br>AGGCCTTCACCCTAGTCATCTCTATGTGGAATACAATTGGGATTCATGAGAGAATCATGCAAGCCA<br>AAGTCAGCAGGTAGCCTTATTTGATGGAGTTAACCAACAGCCAAATTTATTGCCCAGAGCCCATTA<br>ACCATGCTAGAGGAGGGTTTCTCCTGATGACTGGCTCCCTGACATCTGTGTCTGGGCCTCAAATT<br>GGTTTGAAGCTGATGGGCTATGCATAATCCCATCACTAAGACCTTGGTCTTGCTTGCTAGAACTTT<br>TCATTACACCACTAGCTGAAGTTTGTTATGACTGCCCCTAATGGCAGGCTCCTCAGGGACAGAAG<br>GAACTGGCCAACCCCATATCTAATGTTGTTGGGCTTGCTGTAGGGTACCCATCTATGAGGAGGCTT<br>GTAACTGTGATGCTCCCTTAGCTACTTTTGATGTATATGCACAGTGTTGGCAGTGGACCTTGCTCC<br>ATGCAATTCATATGGCTATCTTGCAGTCCATGGCCCACCCCAGACTTCATTGCTGCCCTGTGACAC<br>CATTCATTAGAACCTGGTATGGAAGTGTACCACAGAGCTGCAAGGGGTTGTAGACCACTTATGGT<br>GAAATTTGTAGAATCCAGGGTGGTGAGCTGTAAATGAAACTTTGGATAACTGGCTTTCTAGAGCCT<br>TCCTCCCACCCTCCCCTGTTTGCCATGCCCCAGGGGCAAGCACTGCCCTTTGCACTCTCACTACA<br>GGTAAAGAGTGTCCCCAGGAGTAAATTTCTAGCCCCATAGAAAAGGAAGGTCTAGAGGGAAATTG<br>GCAATGGGCACCTGTCCCATTATAAGCATCTATTTG |
| HIV<br>RRE<br>DNA<br>templat<br>e | GGCTTATCGAAATTAATACGACTCACTATAGGAGCTTTGTTCCTTGGGTTCTTGGGAGCAGCAGGA<br>AGCACTATGGGCGCAGCGTCAATGACGCTGACGGTACAGGCCAGACAATTATTGTCTGATATAGT<br>GCAGCAGCAGAACAATTTGCTGAGGGCTATTGAGGCGCAACAGCATCTGTTGCAACTCACAGTCT<br>GGGGCATCAAACAGCTCCAGGCAAGAATCCTGGCTGTGGAAAGATACCTAAAGGATCAACAGCTC<br>C |
| HIV<br>RRE<br>RNA<br>sequen<br>ce | GGAGCUUUGUUCCUUGGGUUCUUGGGAGCAGCAGGAAGCACUAUGGGCGCAGCGUCAAUGAC<br>GCUGACGGUACAGGCCAGACAAUUAUUGUCUGAUAUAGUGCAGCAGCAGAACAAUUUGCUGAG<br>GGCUAUUGAGGCGCAACAGCAUCUGUUGCAACUCACAGUCUGGGGCAUCAAACAGCUCCAGGC<br>AAGAAUCCUGGCUGUGGAAAGAUACCUAAAGGAUCAACAGCUCC |

**Table S2. Primers used for DNA template amplification.**

| | |
|---|---|
| EGFP_for | GAATTCTAATACGACTCACTATAGGTAGCTAAGG |
| EGFP_rev | CGTATGTTGTGTGGAATTGTGAG |
| 23SdomIIV_for | CTTAAGTAATACGACTCACTATAGCCCTGGCAGTCAGAGG |
| 23SdomIIV_rev | TGAGTCTCGGGTGGAGACAG |
| M13o44_for | TAATACGACTCACTATAGGGTCTCGCTGGTGAAAAGAAA |
| M13o44_rev | AGGCCGGAGACAGTCAAATC |
| rsc1218v1_for | GGCTTATCGAAATTAATACGACTCACTATAGGGCCTAGTGAGTGCATTAGAG |
| rsc1218v1_rT55_rev | GTGGTACACTTCCATACCAGGTTC |
| rsc1218v1_rT77_rev | CAAATAGATGCTTATAATGGGACAGGTGC |
| HIV_RRE_for | GGAGCTTTGTTCCTTGGGTTCTTGG |
| HIV_RRE_T7_for | GGCTTATCGAAATTAATACGACTCACTA |
| HIV_RRE_rev | GGAGCTGTTGATCCTTTAGGTATCTTTC |

**Table S3. Staple sequences used to fold the origami structures.** Named by [object name]_[scaffold name]_[routing variation]_stap[#].

| | |
|---|---|
| rT66_EGFP_Aform_stap1 | GTGGTCGGGGTACAGCTCCTCGCCCTTTTGTGTTGCTGCTTCAT |
| rT66_EGFP_Aform_stap2 | CACTGCACGCCAGGATGGGCACCACCCCGGTGAAGCGGCTGAAG |
| rT66_EGFP_Aform_stap3 | TGGTAGTGGTCGGTTTTTCGAGCTGCACATGGCGGACTTGTTTTTAAGAAGTCGGGAATTGTGAGCGTTTTTGATAACAAT |
| rT66_EGFP_Aform_stap4 | AACAGCTATGAGTCGCCGATGGGGGTGTTCTGCTTCACACAGGA |
| rT66_EGFP_Aform_stap5 | CCAAGCTTGCATGTCGGGCAGCAGCACGGGGCCCCATGATTACG |
| rT66_EGFP_Aform_stap6 | TCCAGCTTACGTTGTGGCTGTTGTAGTCTTTGC |
| rT66_EGFP_Aform_stap7 | TCAGGGCGATCGCGCTTCTCGTTGGGGTTGTAC |
| rT66_EGFP_Aform_stap8 | GTGCCCCAGGACCATGATATAG |
| rT66_EGFP_Aform_stap9 | ATGCCGTTCTTCTTTTTTGCTTGTCGGTGTTGCCGTCCTCTTTTTCTTGAAGTCGGCGCGGGTCTTGTTTTTTAGTTGCCG |
| rT66_EGFP_Aform_stap10 | TCGATGTTGTGGCTCCTGGACGTAGCCTTCGGGCGCTGCCGTCC |
| rT66_EGFP_Aform_stap11 | AGTTCACCTTGTCGTCCTTGAAGAAGATGGTGCGCGGATCTTGA |
| rT66_EGFP_Aform_stap12 | CGAACTCCAGCAGTTTTTGACCATGTGGACTGGGTGCTCATTTTTGGTAGTGGTTGCCTGCAGTTACTTTTTTTGTACAGC |
| rT66_EGFP_Aform_stap13 | CGCTGAACCATCGCCCTCGCCCTCGCTCCCGGC |
| rT66_EGFP_Aform_stap14 | GGCGGTCATCGTCCATGCCGAGAGTGACGGACA |
| rT66_EGFP_Aform_stap15 | TTGTGGCCGTTGCCGTAGGTGG |
| rT66_EGFP_Aform_stap16 | AGATGAACTTCAGTTTTTGGTCAGCTTTACGTCGCCGTCCTTTTTAGCTCGACCGTAGGTCAGGGTGTTTTTGTCACGAGG |
| rT66_EGFP_Aform_stap17 | CGCCCTCGGCTCGATGCGGTTCACCAGCTTGCC |
| rT66_EGFP_Aform_stap18 | GGTGGTGCGTGGGCCAGGGCACGGGCAGGGTGT |
| rT66_EGFP_Aform_stap19 | AACTTCACCTCGATGCCCTTCA |
| rPB66_23s_Aform_stap1 | TTTTCACCCGCTTTTTTTTATCGTTACAACCTTGCTACAGTTTTTAATATAAGT |
| rPB66_23s_Aform_stap2 | GCTTTTCCTGGCTTGGTCTTCCGGCGAGCGGGCGATGCTTAGAG |
| rPB66_23s_Aform_stap3 | TTGTTGCTTCACGATTAACGTTGGACAGGAACCAAGCAGGGCAT |
| rPB66_23s_Aform_stap4 | TTCGCACTTCTCCTTCGGCTCCCCTATTCGGTTTTATGTCAGCA |
| rPB66_23s_Aform_stap5 | CATGCCTCACACGCCCAGTTAAGACTCGGTTTCGATACCTCCAG |
| rPB66_23s_Aform_stap6 | TCAACATTAGTCGTTTTTGTTCGGTCCGGGTCTATACCCTTTTTTGCAACTTAAGCACACCTTCGCATTTTTGGCTTACAG |
| rPB66_23s_Aform_stap7 | TCACAGCACGTGTCCCGCCCTACTCATCATCCG |
| rPB66_23s_Aform_stap8 | CTAATTTTTTTCACCCCCAGCCACAAGTCGAGC |
| rPB66_23s_Aform_stap9 | TGTGCATTTTTACAGGATACCA |

| | |
|---|---|
| rPB66_23s_Aform_stap10 | GGTTCTTTTCGCCTTTTTTTTCCCTCAGTGTCGAAACACATTTTTCTGGGTTTC |
| rPB66_23s_Aform_stap11 | GGCTAGATTTACCCAACCTTCAACCTCACTCCC |
| rPB66_23s_Aform_stap12 | CTCGCCGGACGGTTTCAGGTTCTTTTTGCCCAT |
| rPB66_23s_Aform_stap13 | CACCGGGTTTCTCCAGTTAGTG |
| rPB66_23s_Aform_stap14 | TTCGTGCAGGTACCTGACCACCTGTGTCGGTTTTCATTACGCCA |
| rPB66_23s_Aform_stap15 | CGACAAGGAATCTCAAGCGCCTTGGTATTCTCTCGGAACTTACC |
| rPB66_23s_Aform_stap16 | TCGGGTGGAGACATTTTTGCCTGGCCAGGGGTACGATTTGTTTTTATGTTACCT |
| rPB66_23s_Aform_stap17 | TATACAAAAGGCTCTGACTGCCAGGGCTGAGTCCGCTGACCCAT |
| rPB66_23s_Aform_stap18 | ACGCCTAAGCGAGATTAGCACGTCCTTCATCGCTACGCAGTCAC |
| rPB66_23s_Aform_stap19 | TCACCTTACCGACTTTTTGCTTATCGCTGCTCCCACTGCTTTTTTTGTACGTAC |
| rPB66_23s_Aform_stap20 | AGCTTCGCCCGTTTACCGGGGCTTCGTTATAAC |
| rPB66_23s_Aform_stap21 | GGTTCATACCCATTCGGAAATCGCCGGATCAAG |
| rPB66_23s_Aform_stap22 | TTGCGCTAACCAGTTACGGCCG |
| rPB66_23s_Aform_stap23 | AGGATGGTCCCCCTTTTTCATATTCAGGTGTACGGGGCTGTTTTTTCACCCTGT |
| rPB66_23s_Aform_stap24 | CACTATCGGTCTATGGATTCAGTTAATGATAGTCGGTACTGGTT |
| rPB66_23s_Aform_stap25 | TTTAGCCTTGGCCCCCGGTTCGCCTCATTAACCAGTCAGGAGTA |
| rPB66_23s_Aform_stap26 | TGCCGCAGCTTCGTTTTTGTGCATGGTCTATCTCCCGGTTTTTTTTGATTGGCC |
| rPB66_23s_Aform_stap27 | TAGGGGTCGACTCTTTTTACCCTGCCCGCACCGTAGTGCCTTTTTTCGTCATCA |
| rPB66_23s_Aform_stap28 | ACGCCTTTATATTAACCTGTTTCCCAACGCATA |
| rPB66_23s_Aform_stap29 | AGCGTCGCAACGCTCCCCTACCCAACATCGACT |
| rPB66_23s_Aform_stap30 | CGGCCTCGCCTCAAGTACAGGA |
| rPB66_23s_Aform_stap31 | CCTTCCCAAAGCCAACATCCTGGCTGCAACATA |
| rPB66_23s_Aform_stap32 | GCCTTCTCGGGACAACCGTCGCCCGGCTCTGGG |
| rPB66_23s_Aform_stap33 | CATCGTTTCCCTGGCTGCTTCT |
| rPB66_23s_Aform_stap34 | AACCAGCCTACACTTTTTGCTTAAACCCGTCCCCCCTTCGTTTTTCAGTAACAC |
| rPB66_23s_Aform_stap35 | CTAGTTCCTTCACTTTTTCCGAGTTCTTTCGCTACCTTAGTTTTTGACCGTTATCCATCAATTAACCTTTTTTTCCGGCAC |
| rPB66_23s_Aform_stap36 | AGTTACGGACATATCAGCGTGCCTTCCGGATTT |
| rPB66_23s_Aform_stap37 | GCCTGGAACGCCTCAGCCTTGATTTTCTCCCGA |
| rPB66_23s_Aform_stap38 | CACCATTTTGCTCGCTTCACCT |
| rPB66_23s_Aform_stap39 | TGTGTCTCCCGTGTTTTTATAACATTCAACAGTGCTCTACTTTTTCCCCGGAGA |

| | |
|---|---|
| rPB66_23s_Aform_stap40 | CGGACGTTTCTGGGTTGTTTCCCTCTGCTGGTA |
| rPB66_23s_Aform_stap41 | TCTTCGACTAATAAACAGTTGCAGCCATCACGA |
| rPB66_23s_Aform_stap42 | AGCACCCGCCGTTAGCTGGCGG |
| rPB66_23s_Aform_stap43 | CTATTACGCTTTCTTTTTTTTAAATGAACTTAACCATGACTTTTTTTTGGGACC |
| rPB66_23s_Aform_stap44 | ACATCTTCCGCAATAGCTTTCGGGGAGAACCAGTTAGCCCCGTT |
| rPB66_23s_Aform_stap45 | CGACCAGTGAGTGAATTCACGAGGCGCTACCTAGCAGGCCGACT |
| rPB66_23s_Aform_stap46 | TGTTTGCACAGTGTTTTTCTGTGTTTTTGATTTCAGCTCCTTTTTACGAGCAAG |
| rPB66_23s_Aform_stap47 | CGGTTGATCGCTCGCCGCTACTGGGGATACGTC |
| rPB66_23s_Aform_stap48 | CACTTTCGCGGGCAGGCGTCACACCGTGAATCT |
| rPB66_23s_Aform_stap49 | TTCTTTTCCTCTGCTCCCCGTT |
| rPB66_23s_Aform_stap50 | CACACTGATTCAGTTTTTGCTCTGGGCGGGGTACTTAGATTTTTTGTTTCAGTT |
| rPB66_23s_Aform_stap51 | GGGATGACCAGTTTGCATCGGGTTGGGCTTCCA |
| rPB66_23s_Aform_stap52 | CTAACACAATCGCGCGCCTTTCCAGACTAAGTC |
| rPB66_23s_Aform_stap53 | CCCCTTGCCGATCCGGTATTCG |
| rO44_M13_Aform_stap1 | TGATAAATTAATGTTTTTCCGGAGAGGCAGGTCATTGCCTTTTTTGAGAGTCTG |
| rO44_M13_Aform_stap2 | ACCGTTCTAGCAATATTTAAATTGTAAACGTTATATGATATTCA |
| rO44_M13_Aform_stap3 | GAGACAGTCAAATTTTTTCACCATCAAATATTTTGTTAAATTTTTATTCGCATT |
| rO44_M13_Aform_stap4 | GCCTGGGGTGCTTTTCACCAGTGAGACAGGCCGTAAAGTGTAAA |
| rO44_M13_Aform_stap5 | TTGGGCGCCAGGGTTTTTTGGTTTTTCCTAATGAGTGAGCTTTTTTAACTCACA |
| rO44_M13_Aform_stap6 | AAGGCTATGTAGCTATTTTTGAGAGACGTAGCGGTTTGTCTACA |
| rO44_M13_Aform_stap7 | CCAGCTGGCGAAATTTTTGGGGGATGTGTTTTCCCAGTCATTTTTCGACGTTGT |
| rO44_M13_Aform_stap8 | TCGCTATTACGTTGTTATCCGCTCACAATTCCATGCGGGCCTCT |
| rO44_M13_Aform_stap9 | AACTGTTGGGAAGTTTTTGGCGATCGGCACAACATACGAGTTTTTCCGGAAGCA |
| rO44_M13_Aform_stap10 | AAATCAGCTCACCATTCGCCATTCAGGCTGCGCAAATTTTTGTT |
| rO44_M13_Aform_stap11 | GTGCCGGAAACCATTTTTGGCAAAGCGTTTTTTAACCAATTTTTTAGGAACGCC |
| rO44_M13_Aform_stap12 | ACGCCAGGGCTGCAAGGCGATTAAGTTCTGGCACCGCTTGGGTA |
| rO44_M13_Aform_stap13 | TGAGCGAGTAACATTTTTACCCGTCGGAACAAACGGCGGATTTTTTTGACCGTA |
| rO44_M13_Aform_stap14 | CCTCAGGAAGACAGCTTTCATCAACATTAAATGGACAGTATCGG |
| rO44_M13_Aform_stap15 | TCGCGTCTGGCCTTTTTTTCCTGTAGCTCGCACTCCAGCCTTTTTAGCTTTCCG |

| | |
|---|---|
| rO44_M13_Aform_stap16 | AGCCCCAATGTACCCCGGTTGATAATTAATATCAAAAACAGAAA |
| rO44_M13_Aform_stap17 | TAAAACTAGCATGTTTTTTCAATCATAAAACAGGAAGATTTTTTTGTATAAGCA |
| rO44_M13_Aform_stap18 | GGTAATCGGAGCAAACAAGAGAATCGTGGGATTCTCCGATGAAC |
| rO44_M13_Aform_stap19 | ACGTTGGTGTAGATTTTTTGGGCGCATTCTGCCAGTTTGATTTTTGGGGACGAC |
| rO44_M13_Aform_stap20 | ATTAATGACGGGAAACCTGTCGTGCCGGTCATGGGATAAGCTGC |
| rO44_M13_Aform_stap21 | CGCTCACTGCCCGTTTTTCTTTCCAGTATCGGCCAACGCGTTTTTCGGGGAGAG |
| rO44_M13_Aform_stap22 | ATCATGGTCCGGGTACCGAGCTCGAAGTTGTTAATTGCTTCGTA |
| rO44_M13_Aform_stap23 | GCAGGTCGACTCTTTTTTAGAGGATCCCATAGCTGTTTCCTTTTTTGTGTGAAA |
| rO44_M13_Aform_stap24 | GCATGCCTAAAACGACGGCCAGTGCCTGCACGTAACCGAAGCTT |
| rO66_23s_Aform_stap1 | GCACCGTAGTGCCTTTTTTCGTCATCACCGGGACAACCGTTTTTTCGCCCGGCC |
| rO66_23s_Aform_stap2 | GCTTTTCCTGGTTACTTATGTCAGCATTCGCACGATGCTTAGAG |
| rO66_23s_Aform_stap3 | TTGTTGCTTCAGGGCTTTTCACCCGCTTTATCGAAGCAGGGCAT |
| rO66_23s_Aform_stap4 | GGGGTACGATTTGTTTTTATGTTACCTTTCTGATACCTCCTTTTTAGCATGCCT |
| rO66_23s_Aform_stap5 | TTCCAGACGCTCTCTGACTGCCAGGGCCGGTTTATCGCGCGCCT |
| rO66_23s_Aform_stap6 | ACACACTGATTAGATTAGCACGTCCTTCATCGCTCCACTAACAC |
| rO66_23s_Aform_stap7 | TCACCTTACCGACTTTTTGCTTATCGCCAGGCTCTGGGCTTTTTTGCTCCCCGT |
| rO66_23s_Aform_stap8 | ACCAGCCTTGATTTTCCGGATTTGCCTTATAAC |
| rO66_23s_Aform_stap9 | GGTTCATACCCATTCGGAAATCGCCGGTGGAAA |
| rO66_23s_Aform_stap10 | ACACGCTTAAACGCCTCAGCCT |
| rO66_23s_Aform_stap11 | TCGGTTTCCCTTCTTTTTGGCTCCCCTCGCAGTCACACGCTTTTTCTAAGCGTG |
| rO66_23s_Aform_stap12 | TCTATACCCTGAGCTCACAGCATGTGCATTTTTCGGGTTTCGGG |
| rO66_23s_Aform_stap13 | CCAGTTAAGACACGTGTCCCGCCCTACTCATCGCAACTTAACGC |
| rO66_23s_Aform_stap14 | AACCTGCCCATGGTTTTTCTAGATCACGTGTACGGGGCTGTTTTTTCACCCTGT |
| rO66_23s_Aform_stap15 | TTCGCAGGCTTCAGTTAGTGTTACCCAACCTTCCACAGCACACC |
| rO66_23s_Aform_stap16 | CCCTACCCAACAACATTAGTCGGTTCGGTCCTCACAGAACGCTC |
| rO66_23s_Aform_stap17 | ACAAGTCATCCGCTTTTTTAATTTTTCAACGCATAAGCGTTTTTTCGCTGCCGC |
| rO66_23s_Aform_stap18 | ACCCATTACTTGCTACAGAATATAAGCTTTCAC |
| rO66_23s_Aform_stap19 | CCCCAGCCATCTCCCGGTTTGATTGGCTCGCTG |
| rO66_23s_Aform_stap20 | TACAAAAGGTAATTCGGTTAAC |

| | |
|---|---|
| rO66_23s_Aform_stap21 | CATCCTGGCTGTCTTTTTTGGGCCTTCCCTTAGCTGGCGGTTTTTTCTGGGTTG |
| rO66_23s_Aform_stap22 | CCCCGGAGATGATGATGGCTGCTTCTAAGCCAACAGTGCTCTAC |
| rO66_23s_Aform_stap23 | CGCTACCTAAATGAGCTATTACGCTTTCTTTAAAATTCACGAGG |
| rO66_23s_Aform_stap24 | CCGCGCAGGCCGATTTTTCTCGACCAGTAGCTTTCGGGGATTTTTGAACCAGCT |
| rO66_23s_Aform_stap25 | CCGATTAACCTTAGGGGTCGACTCACGCCCCGT |
| rO66_23s_Aform_stap26 | TACATCTTAGCTTCGGTGCATGGTTTACCTGCC |
| rO66_23s_Aform_stap27 | CGTTGGACAGGTTTCGGCCTCG |
| rO66_23s_Aform_stap28 | CCTGTTTCCCATCTTTTTGACTACGCCAACCCTTGGTCTTTTTTTCCGGCGAGC |
| rO66_23s_Aform_stap29 | CCAAGTACCTCCGTCCCCCCTTCGCAACCATGA |
| rO66_23s_Aform_stap30 | CTTTGGGACCACATCGTTTCCCACTTAGTAACA |
| rO66_23s_Aform_stap31 | AGGAATATTAAAACATAGCCTT |
| rO66_23s_Aform_stap32 | TGTGTCTCCCGTGTTTTTATAACATTCCGGGATGACCCCCTTTTTTTGCCGAAA |
| rO66_23s_Aform_stap33 | GGATTCAGCCCCGGTTCGCCTCATTACGTTAGC |
| rO66_23s_Aform_stap34 | ACCCGCCGTTTCCCTCTTCACGACGGAACCTAT |
| rO66_23s_Aform_stap35 | TTAATGATAGTTGTTTCAGTTC |
| rO66_23s_Aform_stap36 | TCTTTTCCTCGGGTTTTTGTACTTAGAGTGTCGAAACACATTTTTCTGGGTTTC |
| rO66_23s_Aform_stap37 | TAGCCTTGCACTATCGGTCAGTCAGGGAATCTC |
| rO66_23s_Aform_stap38 | GGTTGATTTCGCTCGCCGCTACTGGGGAGTATT |
| rO66_23s_Aform_stap39 | GAGGATGGTCCCGGTACTGGTT |
| rO66_23s_Aform_stap40 | GGTTCTTTTCGCCTTTTTTTTCCCTCACCCCATATTCAGATTTTTCAGGATACC |
| rO66_23s_Aform_stap41 | TTTTCACTTGTACGTACACGGTTTCAATCGGGT |
| rO66_23s_Aform_stap42 | TGGTAAGTTCCGGTATTCGCAGTTTGCGGTTCT |
| rO66_23s_Aform_stap43 | CCCCTCGCCGGCTCCCACTGCT |
| rPB55_23s_Aform_stap1 | CTAAGCCAACATCTTTTTCTGGCTGTCCGGTTTCAGGTTCTTTTTTTTTTCACT |
| rPB55_23s_Aform_stap2 | CCTTGGTCTTCTTTCTTTAAATGATGGCTGCTTTGGACAGGAAC |
| rPB55_23s_Aform_stap3 | CGGCGAGCGGGAGCTATTACGC |
| rPB55_23s_Aform_stap4 | ACATCGTTTCCGCTCCCACTGCTTGTACGTACATGGGCCTTCCC |
| rPB55_23s_Aform_stap5 | CACTTAACCATCGCCTAAGCGT |
| rPB55_23s_Aform_stap6 | CTTCGGCTCCCCTTTTTTATTCGGTTAATACAAAAGGTACTTTTTGCAGTCACAGACTTTGGGACCTTTTTTTAGCTGGCG |

| | |
|---|---|
| rPB55_23s_Aform_stap7 | AGACGCTTGTCACCCTGTATCGCGCGCAAGACT |
| rPB55_23s_Aform_stap8 | CGGTTTCCTGCAACTTAACGCCCAGTTCTTTCC |
| rPB55_23s_Aform_stap9 | CCCATATTCAGACTTTTTAGGATACCAACTGGGTTTCCCCTTTTTATTCGGAAA |
| rPB55_23s_Aform_stap10 | GACCCATTACCTTGCTACAGAATATAATGGAGG |
| rPB55_23s_Aform_stap11 | ATGGTCCCAGTCAGGAGTATTTAGCCTGTCGCT |
| rPB55_23s_Aform_stap12 | TGGTATTCTCTTCGGCCTCGCCTTAGGGGTCGACTCAAGCGCCT |
| rPB55_23s_Aform_stap13 | ACCTGACCACCGACTACGCCTT |
| rPB55_23s_Aform_stap14 | CTAGTTCCTTCACTTTTTCCGAGTTCTCTCACCCTGCCCCTTTTTGATTAACGT |
| rPB55_23s_Aform_stap15 | GGTTCTTTTCGCTCTGACTGCCAGGGCTTTTGCCCCCTCGCCGG |
| rPB55_23s_Aform_stap16 | CCTTTCCCTCATCCTTCATCGC |
| rPB55_23s_Aform_stap17 | ACGCTTATCGCAGTTTTTATTAGCACGCGGTACTGGTTCATTTTTCTATCGGTC |
| rPB55_23s_Aform_stap18 | TTTCCTGGTGATGTTACCTGATGCTTAATATCA |
| rPB55_23s_Aform_stap19 | CCTTACCGTCGCCGGTTATAACGGTTCGAGGCT |
| rPB55_23s_Aform_stap20 | GTGCATTTTTGTGTTTTTTACGGGGCTCCACTAACACACATTTTTCACTGATTC |
| rPB55_23s_Aform_stap21 | CTACTCATCGATAATGATAGTGTGTCGAAACACCGTGTCCCGCC |
| rPB55_23s_Aform_stap22 | GCTCACAGCATATGGATTCAGT |
| rPB55_23s_Aform_stap23 | AGCACCCGCCGTGTTTTTTGTCTCCCGACCGGGTTTCGGGTTTTTTCTATACCC |
| rPB55_23s_Aform_stap24 | GCGCAGGCCGACTTTTTTCGACCAGTGCTTTTCACCCGCTTTTTTTTATCGTTA |
| rPB55_23s_Aform_stap25 | CATCTTCCTTCGGTGCATGGTTTAGCCTCACGA |
| rPB55_23s_Aform_stap26 | CGGACGTTGTCTGGGTTGTTTCCCTCTCCGTTA |
| rPB55_23s_Aform_stap27 | CCGGAGATCCCTTGCCGAAACAGTGCTCCCTAC |
| rPB55_23s_Aform_stap28 | CCAACAACGCAGGCTTACAGAACGCTCCTACCC |
| rPB55_23s_Aform_stap29 | GCATGCCTCACAGTTTTTCACACCTTCGCATAAGCGTCGCTTTTTTGCCGCAGC |
| rPB55_23s_Aform_stap30 | GAATATTAACCTGTTTTTTTTCCCATCTGTGTCGGTTTGGTTTTTGGTACGATTAAGCAGGGCATTTTTTTTGTTGCTTCA |
| rPB55_23s_Aform_stap31 | AAGTACAGTCCGTCCCCCCTTCGCAGTTTCTGA |
| rPB55_23s_Aform_stap32 | TACCTCCACTTATGTCAGCATTCGCACAACACC |
| rPB55_23s_Aform_stap33 | TTGATTGGCCTTTTTTTTCACCCCCAGCGGTTCGGTCCTCTTTTTCAGTTAGTG |
| rPB55_23s_Aform_stap34 | CTCCCGGTATAGCTTTCGGGGAGAACCAACCGG |
| rPB55_23s_Aform_stap35 | GACAACCGAAACCAGCCTACACGCTTAAGCTAT |
| rPB55_23s_Aform_stap36 | GGGTTGGTAAGTCTTTTTGGGATGACCGAATTCACGAGGCTTTTTGCTACCTAA |

| | |
|---|---|
| rPB55_23s_Aform_stap37 | TCCGGTATTCGTCAACCTGCCCATGGCTAGATCTGATAACATTC |
| rPB55_23s_Aform_stap38 | CAGTTTGCATCTTACCCAACCT |
| rPB55_23s_Aform_stap39 | TGATTTTCCGGATTTTTTTGCCTGGATCGCCCGGCCAACTTTTTATAGCCTTC |
| rPB55_23s_Aform_stap40 | TTCAGTTCTCTTTTCCTCGGGGTACTTTCACGC |
| rPB55_23s_Aform_stap41 | CTCAGCCTGCACCGTAGTGCCTCGTCAAGATGT |
| rPB55_23s_Aform_stap42 | ACTGGGGGAATCTTTTTTCGGTTGATTCCCCGGTTCGCCTTTTTTCATTAACCT |
| rPB55_23s_Aform_stap43 | ACATTAGTCCACAAGTCATCCGCTAATGTTCGC |
| rPB55_23s_Aform_stap44 | TCGCCGCTAGGCTCTGGGCTGCTCCCCTTTTCA |
| rT55_rsc1218_Aform_stap1 | CTACCTGCTGATGCACTCACTAGGCCCGTGGTAATCAAATAAGG |
| rT55_rsc1218_Aform_stap2 | CTTTGGCTTGCTGAGTCTCTAA |
| rT55_rsc1218_Aform_stap3 | AGTTCCTTCTGTCTTTTTCCTGAGGAGAATTTGGCTGTTGTTTTTGTTAACTCCCACTTCCATACCATTTTTGGTTCTAAT |
| rT55_rsc1218_Aform_stap4 | CAGGGCAGCAAAACATTAGATATGGGGTTGGCCGAATGGTGTCA |
| rT55_rsc1218_Aform_stap5 | TGAAGTCTGGGGCAAGCCCAAC |
| rT55_rsc1218_Aform_stap6 | GATGGGATTAGCAAGCAAGACCAAGGTACAGTT |
| rT55_rsc1218_Aform_stap7 | ACAAGCCTAAGTAGCTAAGGGAGCATCCTTAGT |
| rT55_rsc1218_Aform_stap8 | CTAGTGGTGTAATTTTTTGAAAAGTTCTATGCATAGCCCATTTTTTCAGCTTCAAGCCAGTCATCAGTTTTTGAGAAACCC |
| rT55_rsc1218_Aform_stap9 | GGGCAGTCATAGGTTAATGGGCTCTGGGCAATACCTGCCATTAG |
| rT55_rsc1218_Aform_stap10 | ACAAACTTCAGTCCTCTAGCAT |
| rT55_rsc1218_Aform_stap11 | CAACACTGTGCATTTTTTATACATCAACCTCATAGATGGGTTTTTTACCCTACAGTGGGCCATGGACTTTTTTGCAAGATA |
| rT55_rsc1218_Aform_stap12 | GTCCCCTCGAGCCCCTCATGTCCTTGGGCAAGG |
| rT55_rsc1218_Aform_stap13 | TCCACTGCGCCATATGAATTGCATGGACCTAGG |
| rT55_rsc1218_Aform_stap14 | CCTGACTAGACATTTTTTTAGTGCATATTCTGGTCATGGATTTTTTACATGGCTATGATTCTCTCATTTTTTGAATCCCAA |
| rT55_rsc1218_Aform_stap15 | TGTCAGGGAACCAATTTGAGGCCCAGAGACTAG |
| rT55_rsc1218_Aform_stap16 | GGTGAAGGTTGTATTCCACATAGAGATCACAGA |
| rT77_rsc1218_Aform_stap1 | AGCCAGTCATCTGCACTCACTAGGCCCCAAATAAGATGTCAGGG |
| rT77_rsc1218_Aform_stap2 | TCCTCTAGCATGATACATGGCTTGAGTCTCTAAAGGAGAAACCC |
| rT77_rsc1218_Aform_stap3 | GGTTAATGGGCTTCTGGTCATG |
| rT77_rsc1218_Aform_stap4 | TGAAGTCTGGGGTTTTTTGGGCCATGGAACCAATTTGAGGTTTTTCCCAGACACGATGCTTATAATGTTTTTGGACAGGTG |
| rT77_rsc1218_Aform_stap5 | TTTCCCTCTAGGAATGGTGTCACAGGGCAGCAACCCATTGCCAA |

| | |
|---|---|
| rT77_rsc1218_Aform_stap6 | CTATGGGGCTACACTTCCATACCAGGTTCTAATACCTTCCTTTT |
| rT77_rsc1218_Aform_stap7 | GAAATTTACTCGCTCTGTGGTA |
| rT77_rsc1218_Aform_stap8 | ACCCTACAGTTACAAGCCTCCTCATAGCCCTGG |
| rT77_rsc1218_Aform_stap9 | ATTCTACATTTCATTTACAGCTCACCAATGGGT |
| rT77_rsc1218_Aform_stap10 | AATTTCACCATGTTATCCAAAG |
| rT77_rsc1218_Aform_stap11 | GCAAGCCCAACGGAGCATCACA |
| rT77_rsc1218_Aform_stap12 | ATATACATCAAAATTTTTGTAGCTAAGAACATTAGATATGTTTTTGGGTTGGCCCTAGTGGTGTAATTTTTTGAAAAGTTC |
| rT77_rsc1218_Aform_stap13 | GCCATATGAATTATGCATAGCCCATCAGCTTCAACTGCAAGATA |
| rT77_rsc1218_Aform_stap14 | AGGTCCACTGCACCAAGGTCTTAGTGATGGGATTGCATGGAGCA |
| rT77_rsc1218_Aform_stap15 | CAACACTGTGCTAGCAAGCAAG |
| rT77_rsc1218_Aform_stap16 | GGAGGAAGGCTCTTTTTTAGAAAGCCAAAGTGGTCTACAATTTTTCCCCTTGCACTGGGGACACTCTTTTTTTTACCTGTA |
| rT77_rsc1218_Aform_stap17 | AGTGCATATAGGGTGAAGGCCTGACTAGGGGCA |
| rT77_rsc1218_Aform_stap18 | TGGCAAACAAGGGCAGTGCTTGCCCCTGACATT |
| rT77_rsc1218_Aform_stap19 | AGGGGAGGGTGGTGAGAGTGCA |
| rT77_rsc1218_Aform_stap20 | GAGCCCCTCATATAGAGATGAC |
| rT77_rsc1218_Aform_stap21 | ATGAATCCCAATTTTTTTGTATTCCACGTCCTTGGCCTAGTTTTTGGTCCCCTCTCTGGGCAATAAATTTTTTTTGGCTGT |
| rT77_rsc1218_Aform_stap22 | CAGTCATATCCCTGAGGAGCCTGCCATTGACTT |
| rT77_rsc1218_Aform_stap23 | TGGCTTGCATCAAATAAGGCTACCTGCTAGGGG |
| rT77_rsc1218_Aform_stap24 | ATGATTCTCTCTGGTTAACTCC |
| rT77_rsc1218_Aform_stap25 | ACAAACTTCAGAGTTCCTTCTG |
| rT66_EGFP_AltAform_st1 | GGTCGGGGTAGAGCTCCTCGCCCTTTTGTGTGGCTGCTTCATGT |
| rT66_EGFP_AltAform_st2 | CTGCACGCCGTGATGGGCACCACCCCGGTGAACCGGCTGAAGCA |
| rT66_EGFP_AltAform_st3 | CAGCTATGACCGTCGCCGATGGGGGTGTTCTGCCACACAGGAAA |
| rT66_EGFP_AltAform_st4 | AAGCTTGCATGTGTCGGGCAGCAGCACGGGGCCATGATTACGCC |
| rT66_EGFP_AltAform_st5 | CCAGCTTGACGTTGTGGCTGTTGTAGTCTTTGC |
| rT66_EGFP_AltAform_st6 | TCAGGGCGTCGCGCTTCTCGTTGGGGTTGTACT |
| rT66_EGFP_AltAform_st7 | TGCCCCAGGATCCATGATATAG |
| rT66_EGFP_AltAform_st8 | TCGATGTTGTGTCCTGGACGTAGCCTTCGGGCACGCTGCCGTCC |
| rT66_EGFP_AltAform_st9 | AGTTCACCTTGGTCCTTGAAGAAGATGGTGCGCGCGGATCTTGA |

| | |
|---|---|
| rT66_EGFP_AltAform_st10 | CTGAACTTATCGCCCTCGCCCTCGCCGCCGGCG |
| rT66_EGFP_AltAform_st11 | GCGGTCACGTCCATGCCGAGAGTGATCGACACG |
| rT66_EGFP_AltAform_st12 | GTGGCCGTTTACCGTAGGTGGC |
| rT66_EGFP_AltAform_st13 | CCCTCGAACTCGATGCGGTTCACCAGGTTGCCG |
| rT66_EGFP_AltAform_st14 | GTGGTGCAGGGCCAGGGCACGGGCAGCGTGTCG |
| rT66_EGFP_AltAform_st15 | CTTCACCTCGGATGCCCTTCAG |
| rT66_EGFP_AltAform_st16 | TGGTAGTGGTCTTTTTGGCGAGCTGCATGGCGGACTTGTTTTTAAGAAGTCGTGAATTGTGAGCGTTTTTGATAACAATTT |
| rT66_EGFP_AltAform_st17 | ATGCCGTTCTTTTTTTCTGCTTGTCGGGTTGCCGTCCTTTTTTCCTTGAAGTCGCGCGGGTCTTGTTTTTTAGTTGCCGTC |
| rT66_EGFP_AltAform_st18 | GAACTCCAGCATTTTTGGACCATGTGAGACTGGGTGCTTTTTTCAGGTAGTGGTCCTGCAGTTACTTTTTTTGTACAGCTC |
| rT66_EGFP_AltAform_st19 | GATGAACTTCATTTTTGGGTCAGCTTGCGTCGCCGTCCTTTTTAGCTCGACCAGAGGTCAGGGTGTTTTTGTCACGAGGGT |
| rT66_EGFP_SymAform_st1 | CGCTCCTGGAACTTGTGGCCGTTTACCCTCGCCG |
| rT66_EGFP_SymAform_st2 | GACACGCTGACGTAGCCTTCGGGCAGATGGTG |
| rT66_EGFP_SymAform_st3 | TGTCGGGCAGCTTTTTTTAGCACGGGGCCGCCGTAGGTGGTTTTTTTCATCGCCCTCGCGTCGCCGTCCTTTTTTTAGCTCGACCAG |
| rT66_EGFP_SymAform_st4 | CGGTGAACGACTGGGTGCTCAGGTAGTGGTGATGGGCACCACCC |
| rT66_EGFP_SymAform_st5 | TTCAGGGTCAGCTTGTCGCCGATGGGGGTGTTCTGCAGATGAAC |
| rT66_EGFP_SymAform_st6 | ATGGCGGACTTGTCCTTGAAGA |
| rT66_EGFP_SymAform_st7 | TTGCTCAGGGCGAGCTCCTCGCCCTTTTGTGTGGGTTGGGGTCT |
| rT66_EGFP_SymAform_st8 | TTGCCGGTGGTGCTGGTAGTGGTCGGCGAGCTGCAACGGGCAGC |
| rT66_EGFP_SymAform_st9 | CGCTGCCGTCCTTTTTTTTCGATGTTGTGTGCTTGTCGGCTTTTTTTCATGATATAGAGGTCACGAGGGTTTTTTTTGGGCCAGGGC |
| rT66_EGFP_SymAform_st10 | CCGTTCTTCGCGGATCTTGAAGTTCATCCCGGCG |
| rT66_EGFP_SymAform_st11 | GCGGTCACTCCATGCCGAGAGTGACCTTGATG |
| rT66_EGFP_SymAform_st12 | GAACTCCAGCATGTACAGCTCG |
| rT66_EGFP_SymAform_st13 | GCTGAAGCCTCCAGCTTGTGCCCCAGGATGTGGTCGGGGTAGCG |
| rT66_EGFP_SymAform_st14 | TTGCCGTCCTCTTTTTTTCTTGAAGTCGACGCGGGTCTTGTTTTTTTTAGTTGCCGTCGAAGAAGTCGTTTTTTTTGCTGCTTCATG |
| rT66_EGFP_SymAform_st15 | CCCTCGAATCGATGCGGTTCACCATGACCATG |
| rT66_EGFP_SymAform_st16 | ATTACGCCACACACACAGGAAACAGCTAGGGTGTCG |
| rT66_EGFP_SymAform_st17 | AGCTTGCATGCTTTTTTTCTGCAGTTACTGGACCATGTGATTTTTTTTCGCGCTTCTCAATTGTGAGCGTTTTTTTGATAACAATTT |
| rT66_EGFP_SymAform_st18 | CTTCACCTCGGTGCCCTTCAGC |
| rT66_EGFP_SymAform_st19 | TAGTTGTAACTGCACGCCGTAGGTCAGGGTCGTTGTGGCTGTTG |

| | |
|---|---|
| rPB66_23S_AltAform_st1 | ACAAAAGGTACCTGACTGCCAGGGCTGAGTCTCTGACCCATTAT |
| rPB66_23S_AltAform_st2 | CCTAAGCGTGCATTAGCACGTCCTTCATCGCCTGCAGTCACACG |
| rPB66_23S_AltAform_st3 | ACCTTACCGACTTTTTGCTTATCGCAGTCCCACTGCTTTTTTTGTACGTACACG |
| rPB66_23S_AltAform_st4 | TTCATATCATTCGGAAATCGCCGGTTAAAGAGC |
| rPB66_23S_AltAform_st5 | TTCGCTTGGTTTACCGGGGCTTCGATCTAACGG |
| rPB66_23S_AltAform_st6 | TTCTTTTCGCCTTTTTTTTCCCTCACGTCGAAACACACTTTTTTGGGTTTCCCC |
| rPB66_23S_AltAform_st7 | CTATCGGTCAGGGATTCAGTTAATGATAGTGTGGTACTGGTTCA |
| rPB66_23S_AltAform_st8 | TAGCCTTGGAGCCGGTTCGCCTCATTAACCTATTCAGGAGTATT |
| rPB66_23S_AltAform_st9 | CACTGATTCAGTTTTTGCTCTGGGCTGGTACTTAGATGTTTTTTTTCAGTTCCC |
| rPB66_23S_AltAform_st10 | TTTTCCTCGGGCTCCCCGTTCG |
| rPB66_23S_AltAform_st11 | TTGATTTCCTCGCCGCTACTGGGGGAACGTCCA |
| rPB66_23S_AltAform_st12 | CTTTCGTGGCAGGCGTCACACCGTATATCTCGG |
| rPB66_23S_AltAform_st13 | AACACACAGCGCGCCTTTCCAGACGCTGTCGGG |
| rPB66_23S_AltAform_st14 | ATGACCCCGTTTGCATCGGGTTGGTAATCCACT |
| rPB66_23S_AltAform_st15 | GATGGTCCCCCTTTTTCATATTCAGACTACGGGGCTGTTTTTTCACCCTGTATC |
| rPB66_23S_AltAform_st16 | GCATTTTTGTGAGGATACCACG |
| rPB66_23S_AltAform_st17 | CAGCATGTTGTCCCGCCCTACTCATCGTCCGCT |
| rPB66_23S_AltAform_st18 | AATTTTTCCACCCCCAGCCACAAGTCAAGCTCA |
| rPB66_23S_AltAform_st19 | CGCCGGGGGTTTCAGGTTCTTTTTCACCATGGC |
| rPB66_23S_AltAform_st20 | TAGATCACACCCAACCTTCAACCTGCCTCCCCT |
| rPB66_23S_AltAform_st21 | TTTCACCCGCTTTTTTTTATCGTTACTCTTGCTACAGATTTTTATATAAGTCGC |
| rPB66_23S_AltAform_st22 | TCGCACTTCTGTCGGCTCCCCTATTCGGTTAACTATGTCAGCAT |
| rPB66_23S_AltAform_st23 | ATGCCTCACAGCCAGTTAAGACTCGGTTTCCCTATACCTCCAGC |
| rPB66_23S_AltAform_st24 | AACATTAGTCGTTTTTGTTCGGTCCTCTCTATACCCTGTTTTTCAACTTAACGCCACACCTTCGCTTTTTAGGCTTACAGA |
| rPB66_23S_AltAform_st25 | CGGGTTTCGGGCAGTTAGTGTT |
| rPB66_23S_AltAform_st26 | TGCCGCAGCTTTTTTTCGGTGCATGGTTCTCCCGGTTTTTTTTGATTGGCCTTT |
| rPB66_23S_AltAform_st27 | ACATCTTCCGCAGCTTTCGGGGAGAACCAGCTATTAGCCCCGTT |
| rPB66_23S_AltAform_st28 | CGACCAGTGAGATTCACGAGGCGCTACCTAAATGCAGGCCGACT |
| rPB66_23S_AltAform_st29 | TGTCTCCCGTGTTTTTATAACATTCTCAGTGCTCTACCTTTTTCCCGGAGATGA |
| rPB66_23S_AltAform_st30 | CTTGCCGAAACCGGTATTCGCA |

| | |
|---|---|
| rPB66_23S_AltAform_st31 | CACCCGCCGTGTAGCTGGCGGT |
| rPB66_23S_AltAform_st32 | GACGTTAGCTGGGTTGTTTCCCTCTTCTGGTAT |
| rPB66_23S_AltAform_st33 | CTTCGACTATAAACAGTTGCAGCCAGCACGACG |
| rPB66_23S_AltAform_st34 | CTATTACGCTTTTTTTTCTTTAAATGACTTAACCATGATTTTTCTTTGGGACCT |
| rPB66_23S_AltAform_st35 | ATCGTTTCCCATGGCTGCTTCT |
| rPB66_23S_AltAform_st36 | CTTCCCACAAGCCAACATCCTGGCTGTAACATA |
| rPB66_23S_AltAform_st37 | GCCTTCTCGGACAACCGTCGCCCGGCCCTGGGC |
| rPB66_23S_AltAform_st38 | AGCGTCGCACGCTCCCCTACCCAACAACGACTA |
| rPB66_23S_AltAform_st39 | CGCCTTTCATATTAACCTGTTTCCCATCGCATA |
| rPB66_23S_AltAform_st40 | TTTTCCTGGAATTGGTCTTCCGGCGAGCGGGCTTGCTTAGAGGC |
| rPB66_23S_AltAform_st41 | GTTGCTTCAGCGATTAACGTTGGACAGGAACCCGCAGGGCATTT |
| rPB66_23S_AltAform_st42 | AGGGGTCGACTTTTTTCACCCTGCCCCACCGTAGTGCCTTTTTTCGTCATCACG |
| rPB66_23S_AltAform_st43 | GGCCTCGCCTTCAAGTACAGGA |
| rPB66_23S_AltAform_st44 | ACCAGCCTACATTTTTCGCTTAAACCGCGTCCCCCCTTTTTTTCGCAGTAACAC |
| rPB66_23S_AltAform_st45 | CCTGGAAACCTCAGCCTTGATTTTCCGCCGAAG |
| rPB66_23S_AltAform_st46 | TTACGGCACATATCAGCGTGCCTTCTCGATTTG |
| rPB66_23S_AltAform_st47 | GGGTGGAGACATTTTTGCCTGGCCATCGGTACGATTTGTTTTTATGTTACCTGA |
| rPB66_23S_AltAform_st48 | CGTGCAGGTCGCTGACCACCTGTGTCGGTTTGGATTACGCCATT |
| rPB66_23S_AltAform_st49 | ACAAGGAATTTCAAGCGCCTTGGTATTCTCTACGAACTTACCCG |
| rPB66_23S_AltAform_st50 | AGTTCCTTCACTTTTTCCGAGTTCTCTCGCTACCTTAGTTTTTGACCGTTATAGTCAATTAACCTTTTTTTCCGGCACCGG |
| rPB66_23S_AltAform_st51 | CCATTTTGCCTCGCTTCACCTA |
| rPB66_23S_AltAform_st52 | TTTGCACAGTGTTTTTCTGTGTTTTTAGATTTCAGCTCTTTTTCACGAGCAAGT |
| rPB66_23S_AltAform_st53 | CGCTAACCCCATTACGGCCGCC |
| rO44_M13_Symform_stap1 | CTGTGTGAAATTTTTTTGTTATCCGCTCAAATCACCATTTTTTCAATATGATAT |
| rO44_M13_Symform_stap2 | CACACAACATACGAAGTGAGACCGGCCGGAGACAGTCACAATTC |
| rO44_M13_Symform_stap3 | GCCGGAAGCATTTTTTAAAGTGTAAAGTTAATTGCGTTTTTTTGCGCTCACTGC |
| rO44_M13_Symform_stap4 | CTGCATTAATGTTTTTAATCGGCCAACCAGGGTGGTTTTTTTTTTCTTTTCACC |
| rO44_M13_Symform_stap5 | CTTTCATCAACTTTTTATTAAATGTGAGGAAGGGCGATTTTTTCGGTGCGGGCC |
| rO44_M13_Symform_stap6 | GCCTCAGGAAGTTTTTATCGCACTCCAAACCAGGCAAATTTTTGCGCCATTCGC |

| | |
|---|---|
| rO44_M13_Symform_stap7 | CTCCGTGGGAATTTTTCAAACGGCGGAATGGGCGCATCTTTTTGTAACCGTGCA |
| rO44_M13_Symform_stap8 | TGCGCAACTGTTGGCGAGTAACAACCCGTCGGATTCATTCAGGC |
| rO44_M13_Symform_stap9 | GCCAGCTGGTAATCATGGTCATAGCTGTTTCTCTTCGCTATTAC |
| rO44_M13_Symform_stap10 | GCTTGCATGCCTTTTTTGCAGGTCGACCCGGGTACCGATTTTTGCTCGAATTCG |
| rO44_M13_Symform_stap11 | CGAAAGGGGGATTTTTTGTGCTGCAAGGGTAACGCCAGTTTTTGGTTTTCCCAG |
| rO44_M13_Symform_stap12 | TGATAAATCGTCTGGCCTTCCTGTAGCCAGTCAACCGTTCTAGC |
| rO44_M13_Symform_stap13 | TTTTGTTAAATTTTTTCAGCTCATTTTGAACGCCATCATTTTTAAAATAATTCG |
| rO44_M13_Symform_stap14 | TAATGCCGGAGTTTTTAGGGTAGCTATTACAAAGGCTATTTTTTCAGGTCATTG |
| rO44_M13_Symform_stap15 | CGTGCCAGCCGCTTTCCAGTCGGGTAGCATTAAAACAAACCTGT |
| rO44_M13_Symform_stap16 | GAGAATCGATGTTTTTAACGGTAATCGGTCAATCATATTTTTTGTACCCCGGTT |
| rO44_M13_Symform_stap17 | AGCCCCAAAAATTTTTCAGGAAGATTGTTTAAATTGTATTTTTAACGTTAATAT |
| rO44_M13_Symform_stap18 | TTGTAAAACAGAAAGATAATCGACGGCCAGTGCCAATCACGACG |
| rO44_M13_Symform_stap19 | CAGTATCGTCTGCCAGTTTGAGGGCAAATATATAAGGACGACGA |
| rO44_M13_Symform_stap20 | CGCATTAAATCCTGAGAGTCTGGAGCAAACAATTTGTTAAAATT |
| rO44_M13_Symform_stap21 | GTGCCGGAGCCAGCTTTCCGGCACAAGTTGGCGATTCGCTTCTG |
| rO44_M13_Symform_stap22 | TTGGGCGCGCGCGGGGAGAGGCGGGAGATCTTTTGATTTGCGTA |
| rO44_M13_Symform_stap23 | TGGTGTAGTTGACCGTAATGGGATCAATAGTTAACAGGTCACGT |
| rO44_M13_Symform_stap24 | AACTCACACCTGGGGTGCCTAATGGGATCCTCTAGAAGTGAGCT |
